# Photosynthetic light harvesting and thylakoid organization in a CRISPR/Cas9 *Arabidopsis thaliana* LHCB1 knockout mutant

**DOI:** 10.1101/2021.12.22.473855

**Authors:** Hamed Sattari Vayghan, Wojciech J. Nawrocki, Christo Schiphorst, Dimitri Tolleter, Chen Hu, Veronique Douet, Gaëtan Glauser, Finazzi Giovanni, Roberta Croce, Emilie Wientjes, Fiamma Longoni

## Abstract

Light absorbed by chlorophylls of photosystem II and I drives oxygenic photosynthesis. Light-harvesting complexes increase the absorption cross-section of these photosystems. Furthermore, these complexes play a central role in photoprotection by dissipating the excess of absorbed light energy in an inducible and regulated fashion. In higher plants, the main light-harvesting complex is the trimeric LHCII.

In this work, we used CRISPR/Cas9 to knockout the five genes encoding LHCB1, which is the major component of the trimeric LHCII. In absence of LHCB1 the accumulation of the other LHCII isoforms was only slightly increased, thereby resulting in chlorophyll loss leading to a pale green phenotype and growth delay. Photosystem II absorption cross-section was smaller while photosystem I absorption cross-section was unaffected. This altered the chlorophyll repartition between the two photosystems, favoring photosystem I excitation. The equilibrium of the photosynthetic electron transport was partially maintained by a lower photosystem I over photosystem II reaction center ratio and by the dephosphorylation of LHCII and photosystem II. Loss of LHCB1 altered the thylakoid structure, with less membrane layers per grana stack and reduced grana width. Stable LHCB1 knock out lines allow characterizing the role of this protein in light harvesting and acclimation and pave the way for future *in vivo* mutational analyses of LHCII.

## Introduction

Light is the source of energy for photosynthetic organisms and is essential for the synthesis of organic molecules in the biosphere. Light excites the chlorophyll molecules of the photosynthetic complexes embedded in the thylakoid membrane and the energy provided allows the photo-oxidation of water by photosystem II (PSII). The released electrons move from PSII through the electron transport chain (ETC) via plastoquinol/plastoquinone (PQ), cytochrome b6f (cytb6f), plastocyanin (Pc), to photosystem I (PSI). Light absorbed by PSI drives the electron transport through ferredoxin to reduce NADP^+^+H^+^ to NADPH. This process is coupled with the generation of a proton gradient between the lumen and the stroma side of the thylakoid membrane, a gradient used by the ATP synthase for the generation of ATP. Light is an inconstant source of energy, and photosynthetic organisms experience shifts in light spectrum and intensity. To cope with these changes and optimize the photosynthetic efficiency, the organization of the pigment-protein complexes must be dynamic (for a recent review, see (Rantala, Rantala, & Aro, 2020)). The dynamism of the electron transport chain is also helped by the non-homogeneous distribution of thylakoid membrane proteins. The stacked domains of the thylakoids, called grana, are enriched in PSII, while other domains, composed of non-appressed membranes, are enriched in PSI and are collectively referred to as stroma lamellae. The stacked and unstacked portions of thylakoids are dynamic and contribute to the acclimation to changes in environmental conditions (for a recent review, see (M. P. Johnson & Wientjes, 2020)).

The larger part of the proteins embedded in the thylakoid membrane are pigment protein complexes that function as light harvesters for the two photosystems. The members of the light harvesting complexes are similar in structure and pigment organization. They can be distinguished, by their interactions with the photosystems and spectroscopic properties, into (i) LHCI, tightly bound to PS I, (ii) the minor antenna complexes LHCB4 (CP29), LHCB5 (CP26) and LHCB6 (CP24) associated with PSII, and (iii) LHCII, which acts as an antenna of both photosystems (P. Bos et al., 2019; Chukhutsina, Liu, Xu, & Croce, 2020; Jansson et al., 1992; Wientjes, van Amerongen, & Croce, 2013). LHCI is composed of protein dimers. In higher plants, the most common PSI-LHCI complex contains two heterodimers of LHCI associated to a monomeric PSI core complex (Mazor, Borovikova, Caspy, & Nelson, 2017). These two dimers are composed of different LHCI isoforms: LHCA1 and LHCA4 form one dimer, while LHCA2 and LHCA3 form the second one (Wientjes & Croce, 2011). Each of these isoforms is encoded by a single gene in Arabidopsis (Klimmek, Sjodin, Noutsos, Leister, & Jansson, 2006). Two additional genes, named *Lhca5* and *Lhca6*, encode close homologue proteins. However, these genes are rarely expressed and their protein products are only found in sub-stoichiometric amounts compared to PSI reaction centers (Klimmek et al., 2006).

LHCII is the most abundant membrane protein in photosynthetic eukaryotes accounting for roughly 30% of the thylakoid membrane proteins and harbors half of the total chlorophyll (Peter & Thornber, 1991). The amino acid sequence and structure of LHCII are highly conserved across plant species, with a 58% sequence identity between the different LHCII isoforms. The LHCII, in higher plants, is composed of homo- and heterotrimers of three similar protein isoforms: LHCB1, LHCB2 and LHCB3 (Caffarri, Croce, Cattivelli, & Bassi, 2004). Trimeric LHCII, and monomeric antennae LHCB4, LHCB5 and LHCB6 play a central role in the structure of the PSII-LHCII supercomplexes. Two classes of LHCII trimers are present in stable PSII-LHCII supercomplexes.

The strongly associated trimers (S-trimers), composed of LHCB1 and LHCB2, interact with the monomeric LHCB4 and LHCB5 isoforms and the inner antenna of PSII. The moderately bound trimers (M-trimers), containing also the LHCB3 isoform, interact only with the monomeric antennae LHCB6 and LHCB4, the latter being connected to the PSII core (For recent reviews, see (Croce & Amerongen, 2020; Xiaowei Pan, Cao, Su, Liu, & Li, 2020)).

An extra pool of LHCII trimers (L-trimers) exists in the thylakoid membranes. These trimers have a loose interaction with PSII and therefore are not consistently isolated with the PSII-LHCII complexes (Dekker & Boekema, 2005). The L-trimers, have been characterized when associated to PSI in the PSI-LHCI-LHCII complex and found to be composed by LHCB1 and LHCB2 (Galka et al., 2012; Kouřil et al., 2005). The LHCII trimers can thus serve both as antennae of PSII and PSI (Wientjes et al., 2013). However the amount of LHCII associated with each photosystem is dynamic and in part regulated via the phosphorylation of a threonine residue located close to the N-terminus of the LHCII protein (Allen, 1992; Wientjes et al., 2013). In particular the phosphorylated N-terminus of LHCB2 is capable to interact with PSI (X. Pan et al., 2018), while LHCB1, in the stable PSI-LHCI-LHCII complex, is mostly non-phosphorylated (A. Crepin & Caffarri, 2015; P. Longoni, Douchi, Cariti, Fucile, & Goldschmidt-Clermont, 2015; Pietrzykowska et al., 2014). Despite this difference between LHCB1 and LHCB2, the STATE TRANSITION 7 (STN7) kinase (Bellafiore, Barneche, Peltier, & Rochaix, 2005) and the PROTEIN PHOSPHATASE 1/THYLAKOID-ASSOCIATED PHOSPHATASE 38 (PPH1/TAP38) (Pribil, Pesaresi, Hertle, Barbato, & Leister, 2010; Shapiguzov et al., 2010) regulate the phosphorylation level of both isoforms (Leoni et al., 2013; P. Longoni et al., 2015). The phosphorylation level of both LHCB1 and LHCB2 is linked to the redox state of the PQ pool present in the ETC. In a simplified model, an increased reduction of PQ activates STN7 and thus induces the phosphorylation of its targets (recently reviewed in (F. P. Longoni & Goldschmidt-Clermont, 2021)). Furthermore, the activity of STN7 is regulated via thioredoxins (Ancin et al., 2019), so that the kinase is inactive when plants are exposed to high light intensity, and by the protein amount, for instance, a prolonged over-excitation of PSI by far-red light leads to degradation of STN7 (Willig, Shapiguzov, Goldschmidt-Clermont, & Rochaix, 2011). While the contribution of LHCB2, and its phosphorylation, to the dynamics of the photosynthetic complexes have been described and appears to be conserved across higher plants, the contribution of phosphorylated LHCB1 is less obvious.

LHCB1 is the most abundant isoform composing the trimeric LHCII, based on proteomic analysis a ratio of 7:4:1 for LHCB1:LHCB2:LHCB3 was calculated for Arabidopsis (Aurelie Crepin & Caffarri, 2018; Galka et al., 2012). In Arabidopsis, there are five genes coding for LHCB1: three genes form a cluster on chromosome 1 (named *Lhcb1.1, Lhcb1.2* and *Lhcb1.3*), with *Lhcb1.1* and *Lhcb1.3* sharing a common bidirectional promoter (Leutwiler, Meyerowitz, & Tobin, 1986). These three genes encode identical mature proteins with few amino acid differences in the sequence of the transit peptide. The remaining two genes (*Lhcb1.4* and *Lhcb1.5*) form a cluster on chromosome 2 and code for slightly different mature proteins characterized by three amino acid substitutions compared to the products of the genes on chromosome 1 (McGrath, Terzaghi, Sridhar, Cashmore, & Pichersky, 1992). Furthermore, the product of *Lhcb1.4* has three extra amino acid substitutions and one deletion in the N-terminal portion of the mature protein that also removes the phosphorylable threonine.

Almost complete depletion of LHCB1 has been previously obtained by transforming Arabidopsis plants with specific artificial miRNA (*Lhcb1amiRNA*) (Pietrzykowska et al., 2014). This knock-down line revealed that this isoform is contributing, via STN7-dependent phosphorylation, to regulate the photosynthetic electron transport upon light quality changes (Pietrzykowska et al., 2014). However, the study on multiple mutants, lacking both STN7 and PPH1/TAP38 revealed that LHCB1 can also be phosphorylated by the STN7 homologue STN8 (P. Longoni, Samol, & Goldschmidt-Clermont, 2019). The analysis of *Lhcb1amiRNA* showed that LHCB1 is required for the accumulation of trimeric LHCII and it is important for Non-Photochemical Quenching (NPQ), a collective definition of mechanisms protecting the photosystems from excessive light irradiance (Reviewed in: (Horton & Ruban, 2005)). Strong reduction of LHCB1, along with LHCB2, was obtained by the introduction of an Lhcb2 antisense gene (Andersson et al., 2003). More recently, by crossing the *Lhcb1amiRNA* with a mutant containing an artificial miRNA targeting LHCB2 coding genes plants without LHCII trimers and with lower NPQ were obtained (Nicol, Nawrocki, & Croce, 2019).

However, there is still no information on the impact of LHCB1 loss on the excitation equilibrium between PSI and PSII, or about the distribution of the photosystems in a “LHCII-depleted” thylakoid membrane. Furthermore, knock-down mutants are prone to changes in gene expression allowing a leaky accumulation of the target protein, which is not the case for a complete knockout. To investigate the role of LHCB1, we used a CRISPR/Cas9 based approach to mutate simultaneously the five genes encoding this protein. The potential of said approach was previously presented, allowing to reduce the LHCB1 protein amount below the detection limit (Ordon et al., 2020). By targeting identical regions shared between the five *Lhcb1* genes, it was possible to use only two synthetic gRNAs to generate stable mutant lines deprived of LHCB1. We describe how the constitutive and complete loss of LHCB1 affects the antenna organization around the photosystems, its phosphorylation and the thylakoid structure.

## Results

### Production of lines deprived of LHCB1

The Arabidopsis genome contains five genes encoding *quasi* identical LHCB1 proteins. These genes are organized in two close regions. Chromosome 1 contains *Lhcb1.1* (AT1G29920), *Lhcb1.2* (AT1G29910) and *Lhcb1.3* (AT1G29930). The remaining two, *Lhcb1.4* (AT2G34430) and *Lhcb1.5* (AT2G34420) are located on chromosome 2. Due to their close proximity it would not be possible to obtain multiple insertional mutants by crossing. A CRISPR/Cas9 approach was used to insert simultaneously a mutation in all the genes. As these genes have largely an identical nucleotide sequence, it was possible to design two gRNAs targeting all of them (Figure 1A). Five plants were transformed with the construct containing the two gRNAs and the gene encoding the Cas9 endonuclease. From each plant, we obtained two to four resistant progenies (T1 generation), which were screened for NPQ, as a loss of *Lhcb1* should lower the photoprotective capacity (Pietrzykowska et al. 2014) (Supplementary Figure S1). The candidates were self-crossed to obtain stable T2 lines. To characterize the nature of the mutation, we sequenced each targeted gene of two independently selected T2 lines. In the line named D2, five different knockout mutations were identified: *Lhcb1.2* (AT1G29910), which has a single nucleotide deletion in the codon for Tyr 191; *Lhcb1.1* (AT1G29920), containing a single nucleotide insertion in the codon for Ser 140; *Lhcb1.3* (AT1G29930), with a single nucleotide insertion in the Pro 192 codon; in *Lhcb1.4* (AT2G34430), a single nucleotide insertion occurred in the codon for Ser 139, and finally *Lhcb1.5* (AT2G34420), harboring a large deletion from the Ser 138 codon to Pro190 codon. A second line, named C2, displayed a deletion/rearrangement between the genes *Lhcb1.2* (AT1G29910) and *Lhcb1.1* (AT1G29920) that made impossible to amplify the gene sequences plus a single nucleotide deletion in the codon for Pro 192 of *Lhcb1.3* (AT1G29930). The mutations on *Lhcb1.4* (AT2G34430), single nucleotide insertion in Ser 139 codon, and *Lhcb1.5* (AT2G34420), single nucleotide insertion in Ser 138 codon, were heterozygous in the T2 line. Therefore, this line was further self-crossed to obtain two T3 lines; one with no mutation in these two genes (C2a) and one carrying the mutations in both genes (C2b). However, even in the presence of non-mutated Lhcb1 genes on chromosome 2, in both C2-derived lines there was no detectable LHCB1 protein nor a detectable band by total protein staining at the LHCII level (Figure 1B). We will refer to the multiple LHCB1 mutants, lacking any detectable LHCB1 protein, as L1ko mutants.

**Figure 1.**
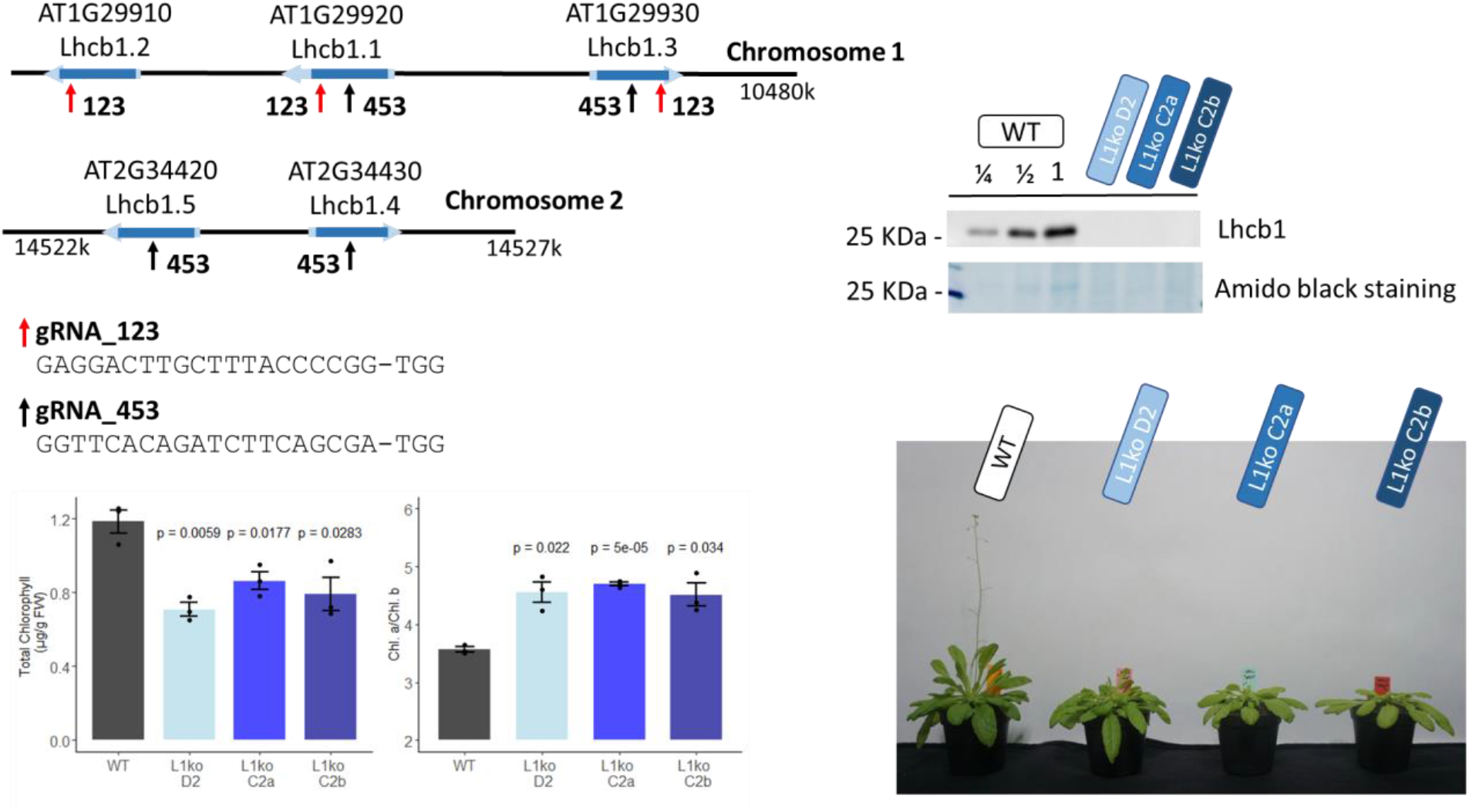
Generation of total knock out lines for the five genes encoding LHCB1. A) Schematic representation of the Lhcb1 gene clusters, showing the regions targeted by the two sgRNAs, the sequence of the sgRNAs is reported below. B) Protein accumulation of the produced mutant lines, the representative immunoblot shows the detection of LHCB1 in a dilution series of WT total protein extract and in the L1ko mutants. Below, the staining of the membrane with amidoblack shows the differential band at 25KDa that corresponds to the LHCII protein C) Total chlorophyll accumulation normalized over the plant fresh weight, left, and chlorophyll a/b ratio, right, in WT and in the three L1ko mutant lines. The error bar indicates the standard error and individual replicates are plotted as points (n=3), above each L1ko line is reported the p value of a Student T-test comparison with WT. D) Representative flowering phenotype of adult plants of the WT and the three independent L1ko lines.

The L1ko mutant plants have a pale green phenotype, with an average chlorophyll per fresh weight content corresponding to 66±6% of the WT (Figure 1C). Consistent with the notion that LHCII is enriched in chlorophyll *b*, this loss was uneven between chlorophyll *a* and *b*, and caused an increase in the chl *a/b* ratio in L1ko lines compared to the WT (Figure 1C). Furthermore, the plant growth was also slower, resulting in a smaller rosette and a delay in flowering (Figure 1D). To compare the generated mutant lines, we analyzed their photosynthetic performance in a range of increasing light intensities. This analysis revealed that there was no statistically significant difference between the three L1ko lines in terms of non-photochemical quenching (NPQ), quantum yield of the photosystem II (ΦPSII) and the fraction of the closed PSII reaction centers (1-qL) (Supplementary Figure S2).

In summary, the CRISPR/Cas9 system based on two sgRNA allowed for the simultaneous mutation of multiple LHCB1-encoding genes. Multiple stable lines produced are characterized by an undetectable amount of the LHCB1 protein and a pale phenotype. All the L1ko lines have a similar photosynthetic activity profile, showing that the loss of LHCB1 correlates with the photosynthetic defect, independently of the presence of non-mutated *Lhcb1.4* and *Lhcb1.5* genes.

### Profile of thylakoid proteins in L1ko

It was previously reported that the removal of LHCB1 by miRNA has a limited impact on the accumulation of other thylakoid proteins, with the exception of LHCB2 (Pietrzykowska et al., 2014). We investigated if the protein accumulation (relative to fresh weight) was modified in the stable L1ko lines compared to WT (Supplementary figure S3). Figure 3 reports the average of the three independent lines D2, C2a and C2b, to highlight differences in protein accumulation due to the loss of the LHCB1 protein, independently of the genetic background. In the L1ko lines, we observed a significant increase in the accumulation of LHCB2 (1.42 ± 0.33) and a small one in LHCB4 (1.27 ± 0.15) compared to WT. There was, however, little to no difference in the accumulation of LHCB5 (1.24 ± 0.40), LHCB6 (1.01 ± 0.19) or LHCB3 (0.89 ± 0.15) (Figure 2). Taken together these results show that the loss of LHCB1 did not cause compensatory changes at the protein level of the minor LHCBs, and confirm the increased accumulation of LHCB2. We also assessed the accumulation of the subunits of the core of PSII (PsbA to D), these proteins did not display any significant difference when compared to WT levels of accumulation (Figure 2).

**Figure 2.**
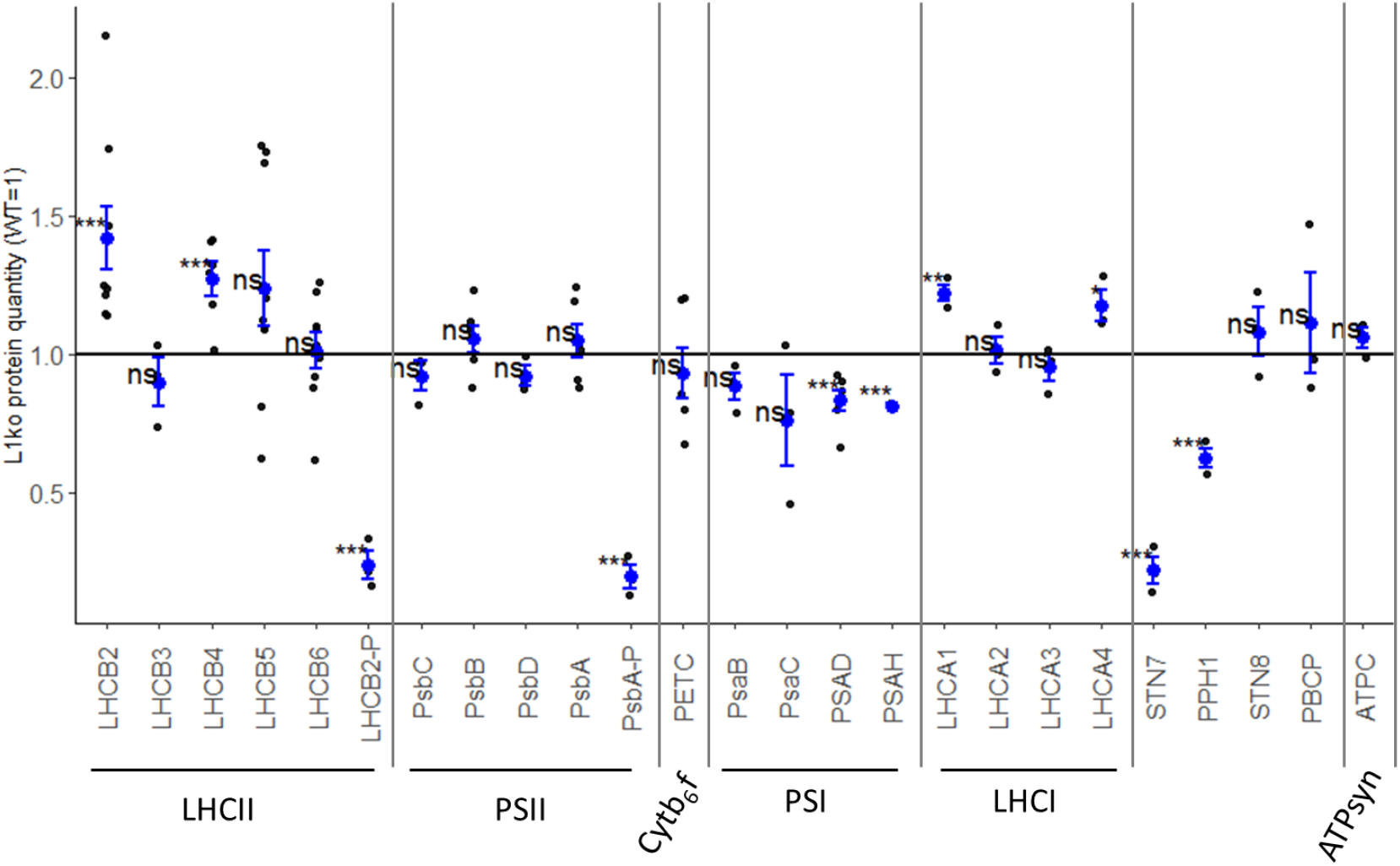
Few thylakoid proteins are affected by the loss of LHCB1. The accumulation of proteins was assessed in 14 days old seedlings of WT and L1ko lines by immunoblotting. The relative protein accumulation, normalized over the actin, was measured compared to the WT level. Each point represent the quantification of the relevant protein in an individual biological replicate, we analyzed one sample per genotype for proteins with low variation between lines and two or three samples for proteins with larger variation. The blue dot represents the average and the blue bars the standard error of the distribution. The protein were divided into the complexes to which they are associated (LHCII, PSII, CytB6f, PSI, LHCI and ATPsyn). The accumulation kinases and phosphatase mainly involved in the regulation of the phosphorylation of LHCII and PSII was also measured. The measured levels were compared to the average of 1, corresponding to the protein accumulation in WT extracts, via a Student’s T-test, asterisks indicate p-value thresholds (0.0001<p<0.001 “***”, 0.001<p<0.05 “**”, 0.05<p<0.10 “*”, 0.10<p “ns”). See also Figure S3a and S3b.

**Figure 3.**
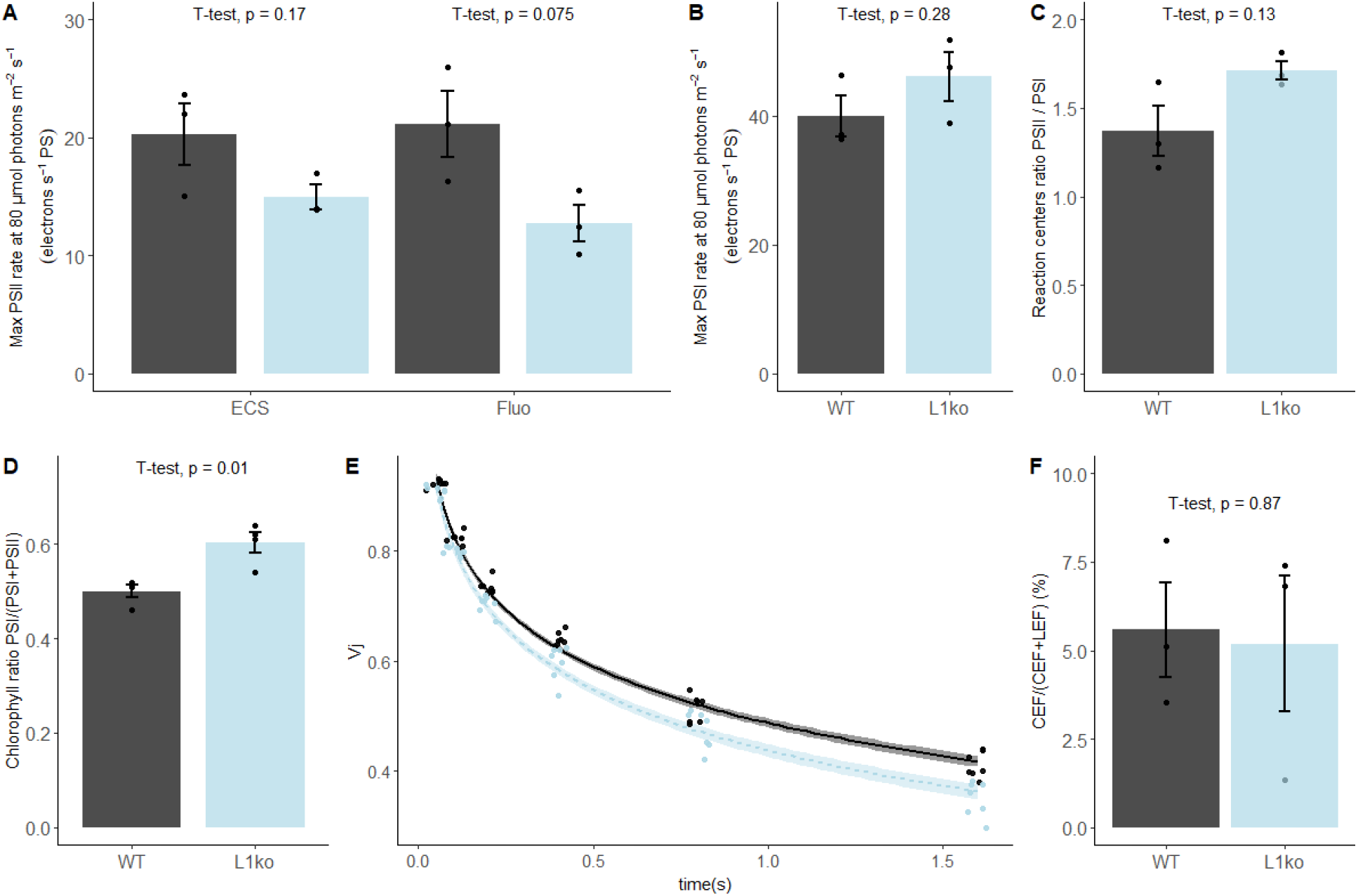
In plants lacking LHCB1, the chlorophyll repartition is shifted towards PSI. A) Maximal photochemical rate of PSII, measured by electrochromic shift (ECS) and fluorescence (fluo) and B) PSI measured by ECS. C) Ratio of PSI/PSII reaction centers measured by a single turnover flash D) Average PSI/(PSI+PSII) chlorophyll ratio determined from the 100ps component of the fluorescence decay measured in fully expanded leaves, excitation wavelength at 633 nm and fluorescence emission at 710-750 nm. (See also Figure S5). E) Fluorescence value measured at 3 ms during a 700 ms saturating pulse normalized over the maximal fluorescence (V_J_), upon sequential incubations of the sample with far red light with increasing time intervals. The points were fitted with a local regression algorithm based on a logarithmic function to visually represent the data distribution in function of the time with a continuous error estimate (black continuous line WT, light blue L1ko). The standard error of the interpolation is shown as a gray area around the curves. F) Contribution of CEF to the total electron flow in WT and L1ko expressed as a percentage of the electron flow. Individual biological replicates are plotted as points, the histogram shows the average and the error bars represent the standard error (n=3, n=4 for panel E). The averages of the distributions were compared with a Student’s T-test, and the p value of the comparison between WT and L1ko reported above the bars.

As for the subunits of PSI, we could not detect a significant difference in the core subunits PsaB and PsaC despite a general tendency towards a lower level of accumulation in L1ko lines compared to WT (0.88 ± 0.09 and 0.76 ± 0.29 respectively). The decrease was instead significant for the peripheral subunits PsaD (0.83 ± 0.09) and PsaH (0.81 ± 0.01), which were less accumulated in L1ko lines compared to the WT (Figure 2). Conversely, two of the four proteins composing the PSI antenna, LHCA1 and LHCA4, were slightly, but significantly, more abundant in L1ko lines compared to WT (1.22 ± 0.06 and 1.17 ± 0.10 respectively), while no significant difference was observed for LHCA2 and LHCA3. We measured the level of PETC protein as a proxy for the cytb6f complex amount; this protein did not show any significant difference in accumulation between L1ko lines and the WT. Similarly, there was no difference in the accumulation level of ATPC, used as a proxy for the ATP synthase complex. The phosphorylation level of LHCB2, and of the two PSII subunits PsbA and PsbC, was clearly lower in L1ko lines compared to the WT (Figure 2, Supplementary Figure S4), as assessed by antibodies recognizing the phosphorylated form of these two proteins, and by phos-tag gels.

Following on the altered phosphorylation of LHCB2 and PSII in L1ko, we assessed the accumulation of the relevant kinases and phosphatases. The accumulation of STN7 (*i.e*. the kinase mainly involved in LHCII phosphorylation) was strikingly lower in the L1ko lines compared to the WT (0.22±0.08). This was also the case for the phosphatase TAP38/PPH1, which counters the activity of STN7, which was less accumulated in L1ko lines compared to the WT (0.62±0.06). Conversely, the second kinase/phosphatase pair, consisting of the thylakoid kinase STN8 and its counteracting phosphatase PBCP, did not show any significant difference in accumulation in L1ko compared to WT.

Taken together these results show that LHCB1 mutation caused a minor change in the composition of the photosynthetic proteins while most of the compensatory responses may occur at the level of their activity and interaction, notably, by the changes in the phosphorylation level.

#### Decrease of PSII antenna size in L1ko is partially compensated by increased PSII/PSI ratio

LHCII antenna has the important role to equilibrate the excitation between the two photosystems. This dynamic equilibrium allows the photosynthetic apparatus to optimize the linear electron transport and to respond to variations in light intensity and quality (Allen, 1992; Ballottari, Girardon, Dall’Osto, & Bassi, 2012). As the L1ko mutants showed an important loss of the LHCII complex (Supplementary Figure S5), we investigated the ratio of the photosystems and their relative antenna size. To assess which proportion of the antenna was connected to the two photosystems in the mutant background the photochemical rate of both photosystems was measured by electrochromic shift (ECS) and fluorescence rise time. This analysis showed that the PSII maximal photochemical rate at 80 μmol photons m^−2^ s^−1^ was 25% lower in L1ko on a per-photosystem basis (14.95 ± 1.78 electrons·s^−1^·PS^−1^) compared to WT (20.25 ± 4.55 electrons·s^−1^·PS^−1^) (Figure 3A), while the PSI maximal photochemical rate was unchanged (WT 39.98 ± 5.48 electrons·s^−1^·PS^−1^, L1ko 46.14 ± 6.56 electrons·s^−1^·PS^−1^) (Figure 3B). The relative amount of the two photosystems partially compensated for the difference in the photochemical rate. The PSII/PSI reaction centers ratio was 1.37 ± 0.25 in the WT and 1.71 ± 0.09 in L1ko (Figure 3C). By multiplying the average photochemical rate measured for PSI and PSII by the average PSII/PSI reaction centers ratio, we can estimate a global PSI/(PSI+PSII) chlorophyll ratio of 0.59 for WT and 0.64 for L1ko. This difference in global chlorophyll ratio between the two photosystems in WT and L1ko was found also by analyzing the fluorescence decay. The calculated PSI/(PSI+PSII) chlorophyll ratio was of 0.52 ± 0.01 for WT and 0.60 ± 0.01 in L1ko (Figure 3D, supplementary Figure S6). The difference between PSI and PSII chlorophylls is also reflected in the rate of Q_B_ oxidation as estimated by the Vj parameter, which is the relative fluorescence measured at 3 ms during a saturating light pulse, normalized over the maximal fluorescence (Toth, Schansker, & Strasser, 2007). In fact, in a leaf exposed to far-red light, the decrease of the Vj value over time is faster in L1ko (Vj trend over time −0.303) compared to the WT (−0.281) (Figure 3E). The same does not occur in the dark, in this situation, the slope of the Vj decrease over time is comparable between the two genotypes: −0.149 for L1ko and −0.147 for WT (supplementary Figure S7). This supports the idea that the faster oxidation of the photoactive plastoquinone is due to the higher activity of PSI over PSII and not to an increased activity of the non-photochemical plastid oxidase (Nawrocki, Tourasse, Taly, Rappaport, & Wollman, 2015). The different distribution in the excitation of the photosystems did not result in a change in the proportion of the two main photosynthetic electron transport routes, in fact the proportion of the cyclic electron flow (CEF) around PSI to the linear electron flow (LEF) was comparable between L1ko and WT (Figure 3F).

The faster PQ oxidation in far-red and the higher PSI/(PSI+PSII) chlorophyll ratio measured in L1ko plants, should result in a larger ΦPSII and open PSII reaction centers under light limiting conditions. To assess that, we performed a follow up experiment using adult plants. The room temperature chlorophyll fluorescence transient on WT and L1ko exposed to a range of increasing light intensities (from 40 to 850 μmol photons m^−2^ s^−1^ PAR) was measured with a fluorescence camera as well as a portable device. Consistently with our hypothesis, the ΦPSII resulted to be higher, especially in the initial range of light intensities used in L1ko compared to WT. The higher quantum yield in L1ko correlated with a larger fraction of open PSII reaction centers estimated by the 1-qL parameter (Figure 4). Furthermore, in agreement with previous results during the screening for L1ko mutants and the literature (Nicol et al., 2019; Pietrzykowska et al., 2014), the NPQ amplitude was lower in L1ko compared to WT (Figure 4). The same trends for NPQ and ΦPSII parameters were confirmed by a second measurement with the portable device (Supplementary Figure S8).

**Figure 4.**
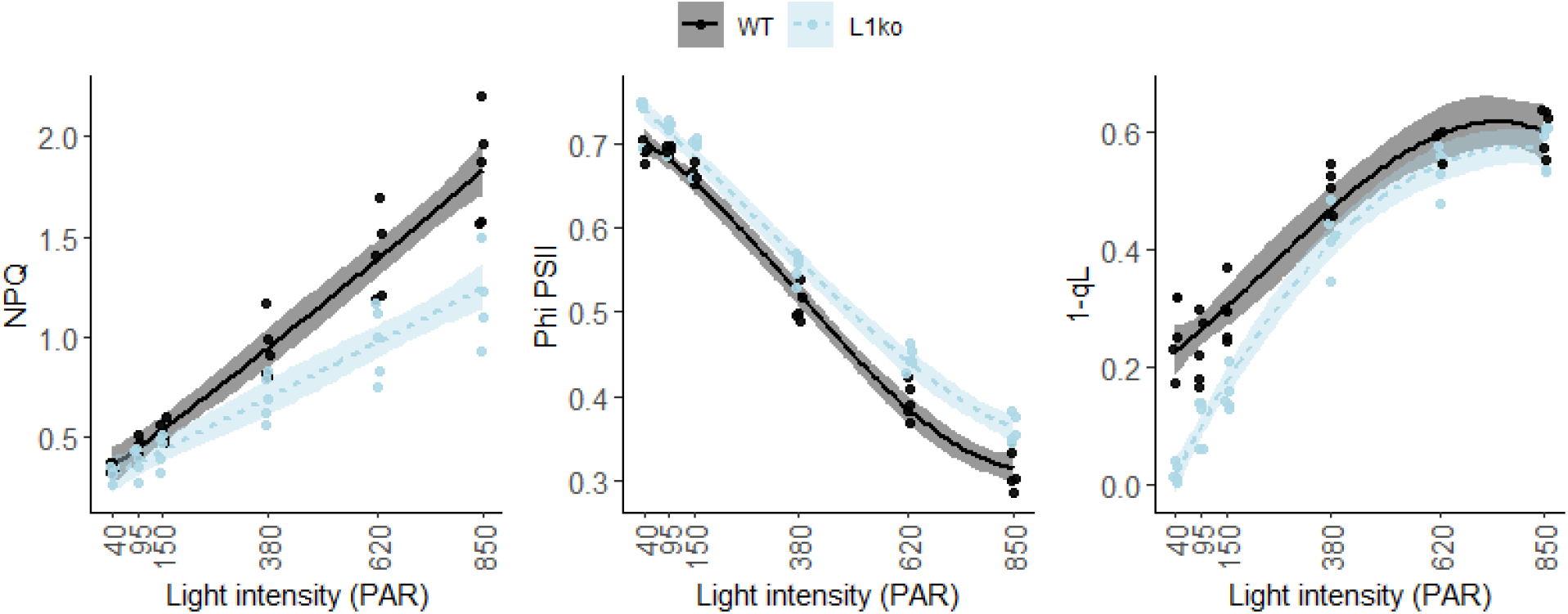
Loss of LHCB1 alters the equilibrium of the LEF between PSII and PSI. The LEF was monitored by room temperature chlorophyll fluorescence on adult WT (black) and L1ko (light blue) plants. The plant were exposed to a range of increasing light intensities, each lasting 5 minutes. A) Non photochemical quenching (NPQ), B) Quantum efficiency of PSII, referred as ΦPSII in the main text. C) Fraction of closed PSII centers estimated by the fluorecence by the 1-qL parameter. Each experimental point is displayed as a dot. The measured points were fitted with a second degree polynomial function for NPQ, a third degree polynomial for φPSII and 1-qL to visually represent the data distribution and the standard error for WT (Black line and gray area) and L1ko (Blue dotted line and light blue area). Post-hoc analysis of the models show that there is a significant difference between L1ko and WT (p<0.001).

In conclusion, the loss of LHCB1 has a larger impact on the photochemical rate of PSII than on that of PSI. This imbalance is partly compensated by an increase of the relative PSII/PSI reaction centers ratio. However, this does not allow L1ko plants to reach the same photosynthetic equilibrium of the WT.

### Loss of LHCB1 alters the thylakoid organization

The trimeric LHCII is a large component of the thylakoid membrane. Consequently, a change in the shape and organization of the thylakoid membrane is expected upon removing LHCB1, which constitutes a large portion of LHCII. Therefore, the chloroplast ultrastructure was investigated in the mutant lines by transmission electron microscopy (TEM) and confocal microscopy. While confocal microscopy allows to observe the overall chloroplasts topology and the distribution of the photosystems, the TEM images allow to observe the fine structure of the thylakoids (Figure 5). Confocal microscopy images of isolated chloroplasts were acquired using two detection ranges to preferentially detect either the emission of PSII (650-680 nm, green), or of PSI (710-750 nm, red) (Figure 5A, Supplementary Figure S9). These images showed relatively more chlorophyll connected to PSI and smaller grana in L1ko compared to the WT (yellow areas in Figure 5A, Supplementary Figure S10). Since the spatial resolution of confocal microscopy is too low, TEM images were used to measure the grana widths and number of grana layers per stack (Figure 5B, Supplementary Figure S11). These analyses showed that the loss of LHCB1 leads to fewer stacks per grana on average (3.75) compared to the WT (4.57). Covariance analysis showed a significant negative correlation between the number of grana and the number of stacks for both the WT and L1ko (z value = −13.8, p < 0.001), but such correlation was different across genotypes (interaction term: z value = 3.12, p = 0.002), which results on an average of 21% more stacks per grana in WT compared to the L1ko mutant (genotype effect; z value = −3.4, p = 0.001). Concerning grana width, the variance analysis shows a significant difference between the genotypes (p < 0.001), with an average grana width for the WT of 500 nm and an average width of 356 nm for the L1ko mutant. Consistently with the observation of the reduced grana width and stacks number, upon separation of the grana stacks from total thylakoids by digitonin, the grana fraction from WT thylakoids contained 66±4% of the total chlorophyll, while in L1ko this percentage was reduced to 40±14% (Supplementary Figure S12).

**Figure 5.**
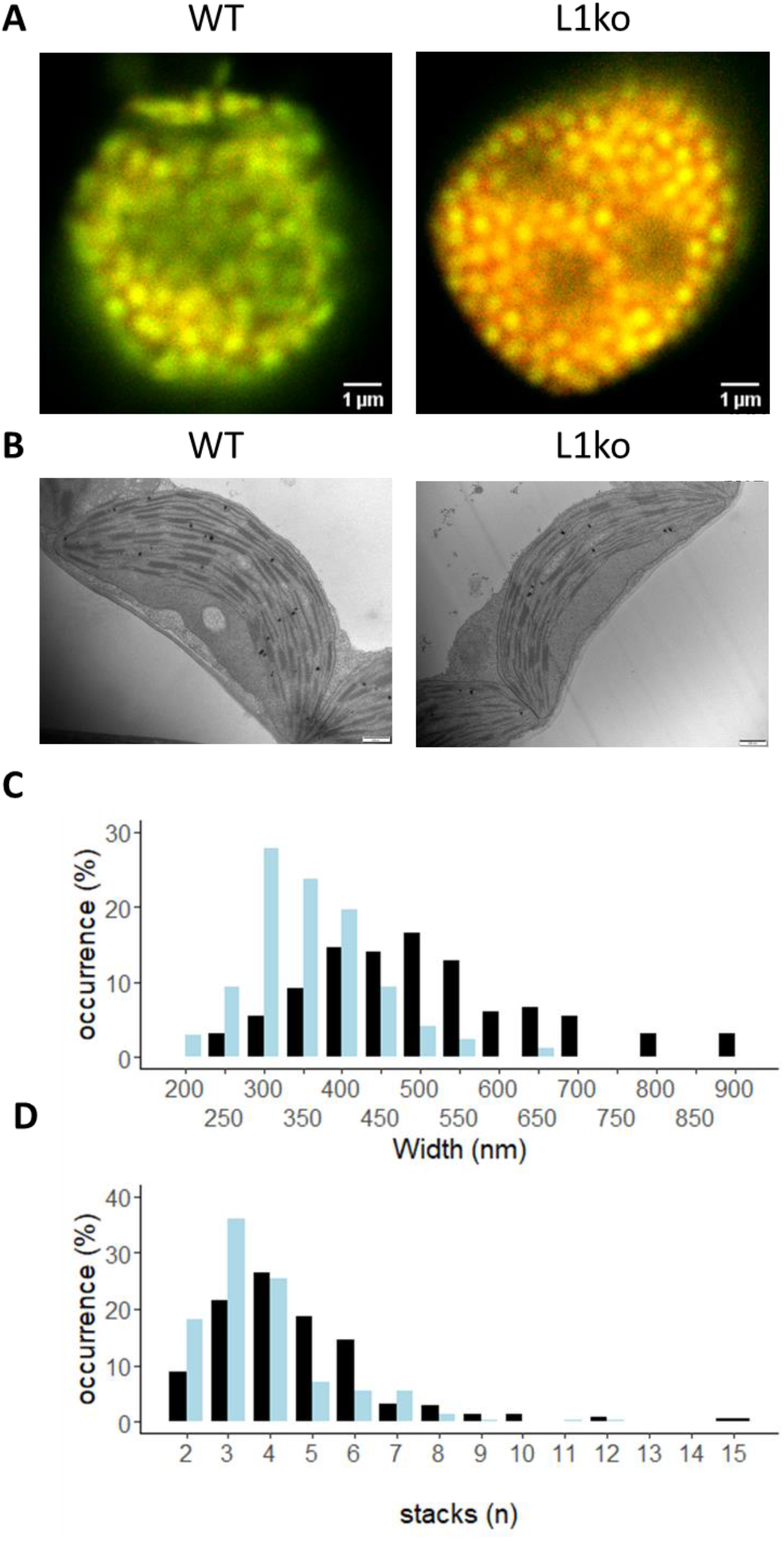
LHCB1 mutation affects the thylakoid structure. A) Confocal microscopy on isolated chloroplasts of WT and L1ko. Excitation at 633nm and detection at 650-680nm is showed in green (PSII-maximum) while 710-750nm detection in red (PSI-maximum). Red channel intensity has been increased 10x in comparison to the green channel to compensate for decreased sensitivity of the detector and decreased fluorescence emission in the far-red region. See also Figure S9. B) Representative images of chloroplasts from WT and L1ko leaves obtained by transmission electron microscopy. The scale bar is 500 nm. See also Figure S11 C) Distribution of the grana width in WT and L1ko based on the measurement of 164 and 173 grana for WT and L1ko, respectively. D) Distribution of the grana stacks divided by number of layers based on the analysis of 215 and 241 grana for WT and L1ko, respectively. The chloroplast images used for the analysis were obtained from two independent biological samples. The statistical analysis of the grana stack number and width distributions is presented in the text.

To assess whenever the change in the LHCII content had an impact on the lipid portion of the thylakoid membrane we measured the total content of the two main classes of galactolipids, monogalactosyl diacylglycerols (MGDGs) and digalactosyl diacylglycerols (DGDGs). This analysis showed that the total galactolipid content, normalized over the fresh weight, was unchanged between the L1ko lines and the WT. In addition, the relative abundance of the different molecules sorted by the degree of saturation and length of the acyl chains was comparable between the L1ko lines and the WT (Supplementary Figure S13). The main prenyl lipids found in the thylakoid membrane were also accumulated to the same level in L1ko and WT, only the xanthophylls, which are mostly associated to LHCII proteins (i.e. Violaxanthin, Neoxanthin, Zeaxanthin and Lutein) were clearly less abundant in the L1ko lines compared to the WT. Conversely, there was no change in β-carotene content (Supplementary Figure S13).

## Discussion

### Three of the five genes encoding LHCB1 are the major contributors to the protein accumulation

Stable multiple mutant lines are fundamental to study the function and role of the major components of the LHCII. The two major isoforms, LHCB1 and LHCB2, are coded by multiple genes, therefore, the production of these mutant lines requires an approach targeting simultaneously multiple genes. The CRISPR/Cas9 technique provided such a tool, allowing the mutation of entire gene families with a single transformation event by using multiple gRNAs along with the Cas9 endonuclease (Ordon et al., 2020). In this report, we targeted conserved sequences, shared by multiple LHCB1-coding genes, to design two gRNAs capable of knocking-down the five genes. The pale green phenotype and the previously reported lower NPQ level in plants in which LHCB1 was depleted by artificial miRNA (*amiLhcb1*) (Pietrzykowska et al., 2014) facilitated the screening procedure allowing a rapid identification of multiple mutant lines characterized by a complete loss of LHCB1. The selected mutants were all mutated in chromosome 1 genes (therefore lacking the Lhcb1.1, Lhcb1.2 and Lhcb1.3 *genes*). Conversely, mutations in *Lhcb1.4* and *Lhcb1.5* were not present in all the lines even if there was no detectable LHCB1 protein. However, it has to be noted that the primary amino acid sequence of the *Lhcb1.4* product is slightly different from the other LHCB1 protein isoforms. This difference, present at the level of the N-terminal portion of the mature protein, may affect the antibody binding, and lead to an under estimation of the *Lhcb1.4* product (Jansson, 1999). However, the line containing a non-mutated version of the *Lhcb1.4* gene along with *Lhcb1.5*, named L1ko “C2a”, did not show any significant difference in chlorophyll content or NPQ when compared to the other L1ko lines (Supplementary Figure S2). Therefore we can conclude that, despite being reported as relatively highly expressed genes (Jansson, 1999), the contribution of *Lhcb1.4* and *Lhcb1.5* to the total accumulation of the LHCB1 protein seems negligible in our growth condition.

### Change in protein phosphorylation and photosystem stoichiometry partially compensate for the loss of LHCB1

Being the most abundant isoform, in L1ko lines the loss of LHCB1 results in the almost complete absence of the trimeric LHCII complex. However, only the accumulation of LHCB2 increased while the LHCB3 level showed no significant difference between L1ko and WT. These moderate changes cannot compensate for the LHCB1 loss and align well with the *amiLhcb1* line phenotype (Pietrzykowska et al., 2014). Except for LHCB4, the accumulation of which increased by around 20% in L1ko compared to WT, the monomeric antennae also show no significant increase. A previous report suggested that the loss of trimeric LHCII was compensated by an increase in the LHCB5 protein (Ruban et al., 2003). LHCB5 was proposed to be integrated into the trimeric LHCII thanks to the presence of an N-proximal trimerisation motif (Hobe, Foerster, Klingler, & Paulsen, 1995). However, a higher accumulation of LHCB5 does not seem to be induced in the L1ko lines as well as in the *amiLhcb1* line (Pietrzykowska et al., 2014) and in the noLHCII mutant (Nicol et al., 2019). Overall, these observations suggest that LHCB5 has no specific role, nor that the accumulation of a specific antenna isoform is induced to compensate the loss of the isoforms composing the LHCII trimers.

Despite similarities between the L1ko mutation via CRISPR/Cas9 and the *amiLhcb1* line, differences exist between the two genotypes, notably at the level of STN7 accumulation. In fact, in the *amiLhcb1* line the amount of the STN7 protein was found to be higher compared to WT (Pietrzykowska et al., 2014). In the mutant lines produced via CRISPR/Cas9 we clearly observed the opposite response (Figure 2). The decrease of STN7 in the L1ko compared to WT is consistent with the previously reported long-term regulation of STN7 abundance in response to changes of light conditions. Exposure of plants to far red light, a condition that enhances excitation of PSI over PSII, leads to a decrease in the amount of the STN7 kinase (Willig et al., 2011). Therefore, the decrease of STN7 could be a consequence of the oxidation of the photosynthetic electron transport chain, as in the L1ko line the chlorophyll distribution is shifted in favor of PSI (Figures 3 and 4). Conversely, the lower level of TAP38/PPH1 observed in the L1ko lines is less obvious. In previous reports the protein level of STN7 and TAP38/PPH1 were shown to be oppositely regulated; *i.e*. a decrease of the STN7 kinase was coupled with an increase of the TAP38/PPH1 phosphatase (Wunder et al., 2013). The observation that both the STN7 kinase and the counteracting phosphatase are less accumulated in the L1ko lines suggests the existence of another layer of regulation of the protein accumulation, besides the previously documented redox status of the photosynthetic ETC. Consistent with this notion, a reduction of TAP38/PPH1 level was also observed in *amiLhcb1* lines (although to a lesser extent) (Pietrzykowska et al., 2014), suggesting that TAP38/PPH1 abundance may be also regulated by the presence of its substrate LHCB1 (Suorsa et al., 2016). The observation that the phosphorylation level of the thylakoid protein was lower in L1ko lines is in agreement with previous reports linking the thylakoid protein phosphorylation to the redox state of the photosynthetic PQ pool more than to the accumulation of the relevant kinases and phosphatases (Suorsa et al., 2016). This appeared to be particularly true for the PSII kinase STN8, as PSBA and PSBC, which are the main targets of this kinase, showed a strongly decreased phosphorylation in the L1ko lines compared to the WT. However, the lower phosphorylation level was not coupled with a decrease of the relevant kinase, STN8, or the PBCP phosphatase. This supports the hypothesis that their activity is regulated in response to changes in the redox status, previously formulated based on the observation of plants overexpressing the STN8 kinase (Wunder et al., 2013).

### Photosynthetic electron transport in L1ko

The clear reduction in STN7 accumulation and phosphorylation of the main thylakoid phospho-proteins in L1ko compared to WT, may be related to an imbalance in the redox equilibrium of the photosynthetic ETC. We therefore investigated the status of the ETC using chlorophyll fluorescence and transient absorption spectroscopy to detail the impact of the LHCB1 loss on the photosynthetic activity. The first expected impact of the absence of LHCB1, and thus a decrease of LHCII trimers, is a reduction of the PSII physical and functional antenna size (Terao & Katoh, 1996). Consistently, both chlorophyll fluorescence and ECS measurements confirmed that in L1ko the maximal PSII photochemical rate measured at a light-limiting intensity of 80 μmol photons m^−2^ s^−1^, was lower in L1ko compared to WT (Figure 3A). Besides the interaction with PSII, several reports have shown that LHCII also contributes to the excitation of PSI (I. Bos et al., 2017; Chukhutsina et al., 2020; Wientjes et al., 2013). Therefore, we used ECS-based methods to investigate whether the absence of LHCB1 affects the antenna size of PSI. Our data showed that the maximal PSI rate at 80 μmol photons m^−2^ s^−1^ in the L1ko is comparable, with a tendency to be larger than that in the WT (Figure 3B). Therefore, no major difference in PSI antenna size should be present between these two genotypes. This can be explained by the relative increase of the LHCA1 and LHCA4 proteins, part of LHCI antenna, over the PSI core (Figure 2). Increased LHCI could compensate for the loss of LHCII preserving the PSI antenna size. This hypothesis fits well with a previous report showing that an extra LHCI dimer, composed by LHCA1 and LHCA4, can functionally connect to PSI (A. Crepin, Kucerova, Kosta, Durand, & Caffarri, 2020). We further defined if there was an excitation imbalance between photosystems in L1ko by an indirect measurement of the rate of PQ oxidation by far red light based on chlorophyll fluorescence rise (Vj) (Pralon et al., 2020; Toth et al., 2007). The result showed that PQ oxidation is faster in L1ko, supporting the idea that the difference between PSI and PSII excitation in far red light is larger in L1ko compared to the WT background (Figure 3E). The same analysis, conducted in the dark, showed no difference, suggesting that the difference in the PQ oxidation between L1ko and WT is due to a different relative excitation of PSI and not to an increased contribution of the non-photochemical oxidation of the PQ pool (Supplementary Figure S7) (Nawrocki et al., 2015). Consistent with this result, the analysis of the fluorescence decay shows a bigger contribution of the faster component, which is interpreted as the decay of PSI excitation (Wientjes, Philippi, Borst, & van Amerongen, 2017). This confirms that, in L1ko, the global relative chlorophyll pool connected to PSI over PSII is larger compared to the WT, therefore the decrease in PSII antenna size is not fully compensated by the change in PSII/PSI reaction center ratio. The increase in the PSII/PSI reaction center ratio is a compensatory response previously observed in chlorophyll *b* mutants as well as in no-LHCII mutants (Nicol et al., 2019; Terao & Katoh, 1996). In terms of maintaining the equilibrium of the PQ pool redox state, the excess excitation of PSI could have been partially compensated by an increase of the CEF/LEF ratio. However, there was no significant difference in the CEF over LEF ratio between the L1ko and WT, suggesting that the CEF/LEF ratio does not play a major role in balancing the ETC redox equilibrium in L1ko (Figure 3F). As the relative PSI/PSII chlorophyll ratio is higher in L1ko than the WT this should lead to higher oxidation of the photoactive PQ pool under light-limiting condition in the mutant. The oxidation of the photoactive PQ can be inferred by a smaller fraction of closed PSII reaction centers (Krause & Weis, 1984). Consistently, when L1ko plants were analyzed by room temperature fluorescence, we observed a smaller fraction of closed PSII centers as determined by the 1-qL parameter (Kramer, Johnson, Kiirats, & Edwards, 2004), and a higher φPSII compared to the WT (Figure 4, Supplementary Figure S8). As discussed in the previous paragraph the constitutive over-oxidation of the PQ pool, could explain the low phosphorylation of the thylakoid proteins, and the low level of STN7 accumulation in L1ko. Furthermore, STN7 activity has been also shown to be important in regulating the grana stacks diameter (Hepworth et al., 2021). With this regard, the low level and activity of STN7 can also have a role in compensating the smaller grana observed in L1ko and contribute to maintain the equilibrium between LEF and CEF in this mutant as proposed in (Hepworth et al., 2021).

### Lack of the LHCB1 isoform alters the thylakoid organization

The trimeric LHCII occupies a large part of the thylakoid membrane surface (Kirchhoff, Mukherjee, & Galla, 2002). Therefore, it has been proposed to play a major role in the structure of the thylakoids and in particular in the grana stacking (Albanese, Tamara, Saracco, Scheltema, & Pagliano, 2020; Barber, 1980; Daum, Nicastro, Austin, McIntosh, & Kühlbrandt, 2010). However, plants deprived of the two major isoforms, LHCB1 and LHCB2, still have grana stacks (Andersson et al., 2003; Nicol et al., 2019; Ruban et al., 2003). The grana were present also in the *amiLhcb1*, but with less grana stacks on average when compared to the WT (Pietrzykowska et al., 2014). Our investigation by TEM found that L1ko contains statistically less stacks per grana compared to WT, and on average a smaller diameter (Figure 5). This was in part expected as, by losing LHCII, the PSII particles in the grana would be smaller and packed more tightly and thus occupy a smaller area (Goral et al., 2012). LHCII phosphorylation by STN7 was linked to the regulation of the grana diameter as discussed earlier (Hepworth et al., 2021). However also the STN8/PBCP kinase and phosphatase were proposed to play a role in regulating grana dynamics. In particular, the mutation of STN8, the kinase mainly involved in the phosphorylation of PSII core protein, was associated with an increase in the grana diameter (Fristedt et al., 2009). Further studies have shown that the CURVATURE 1 (CURT1) protein, which is also phosphorylated in a STN8-dependent fashion, is directly involved in regulating the grana diameter and may be responsible for the change of grana width in the *stn8* mutant (Armbruster et al., 2013; Trotta et al., 2019). Therefore, as for STN7, also the low level of STN8 activity may have a compensatory function in terms of grana width and stability in the L1ko background.

Although the relationship between the protein composition of the thylakoid membrane and the grana structure has been described in detail, there is a less clear consensus about the role played by the membrane lipids in shaping the thylakoid membrane. In this report, we analyzed the galactolipids present in L1ko, and we found no significant difference with the WT (Supplementary Figure S13). This change may affect not only the ultrastructure of the thylakoids, but also the electron transport routes by interfering with the lipid mobility and thus hampering the PQ diffusion between PSII and the cytb_6_f antagonizing the acclimation strategies of L1ko discussed in the previous paragraph (Matthew P. Johnson, Vasilev, Olsen, & Hunter, 2014; Tyutereva, Evkaikina, Ivanova, & Voitsekhovskaja, 2017). Quantification of the carotenoids revealed that, in the L1ko lines, the lutein and violaxanthin/neoxanthin content was lower than in the WT (Supplementary Figure S13). This is not surprising since the large majority of the xanthophylls are expected to be bound to LHCII, while most of the *β*-carotene, which is not decreasing in L1ko compared to WT, is associated to the photosystems core, and mostly to PSI (Qin, Suga, Kuang, & Shen, 2015). The same decrease in xanthophyll content was not observed in the *asLhcb2* line, which is lacking both LHCB1 and LHCB2 proteins (Andersson et al., 2003), while the carotenoid content decreased to the same extent in the crossed *amiLhcb1/amiLhcb2* line (Nicol et al., 2019). Hinting that, in these lines, a signaling mechanism is in place to coordinate the carotenoid metabolism with the antenna biosynthesis (Li et al., 2009).

In conclusion, we report that the stable loss of LHCB1 induces compensatory mechanisms in the plant. These mechanisms include the alteration of the kinase and phosphatases, regulating the equilibrium of the photosynthetic ETC. Taken together these data show that loss of LHCB1 cannot be compensated by the other LHCII isoforms and that this isoform, by its nature or level of accumulation, has a unique role in shaping the thylakoid membrane and maintaining the equilibrium between the two photosystems. Besides, this mutant gives the first possibility for a mutational analysis on the major LHCII, something that up until now was impossible due to the clustered gene organization. Since trimeric LHCII is essential for the plasticity of the photosynthetic machinery that allows the rapid acclimation to changing light conditions, these mutants are not only of vital importance for fundamental research but also for applied projects aiming at increasing crop productivity by improving the light use efficiency.

## Methods

### Plant material and mutant production

For the production of multiple mutant lines two synthetic gRNA (sgRNA) were designed based on the Arabidopsis genomic sequence, using the software chop chop for the design (Montague, Cruz, Gagnon, Church, & Valen, 2014). The highest ranking sgRNA sequences, targeting multiple genes, were further analyzed with E-CRISP (Heigwer, Kerr, & Boutros, 2014). The sgRNA sequences with the highest efficacy score were cloned in a binary vector for the stable transformation of wild type Arabidopsis plants (ecotype: Col0) containing the Cas9 gene under the control of the synthetic EC1 promoter that confers expression in egg cells (Durr, Papareddy, Nakajima, & Gutierrez-Marcos, 2018). The insertion of two sgRNA was performed following the method described by Xing and coworkers (Xing et al., 2014).

Seed obtained from the transformed plants were germinated on ½ MS containing 33 ug ml^−1^ Hygromycin. After 10 days resistant plants were transferred to soil. The genomic DNA was extracted from a leaf sample and a PCR was perform to confirm the presence of the T-DNA insert. Plants containing the T-DNA were lead to flowering and the collected seeds germinated on ½ MS, the seedlings were screened by chlorophyll fluorescence measurement (detailed below) to detect low NPQ individuals within mixed populations (Supplementary Figure 1). Those individuals were transferred to soil and further analyzed for presence of the T-DNA insert and for the presence of the mutation in the Lhcb1.1-5 genes.

The plants on plates were grown under white light LEDs (120 μmol photons m^−2^ s^−1^ PAR) with a 16h light 8h dark daily cycle in a climatic chamber set at 22°C (ARALAB FitoClima 600 PL). The plants in soil were grown under white neon tubes (120 μmol photons m^−2^ s^−1^ PAR) with a 16h light 8h dark daily cycle in a walk-in climatic chamber set at 22°C. The chlorophyll extraction was performed on samples composed of three 14 days-old plantlets, which were weighted to measure the total fresh weight, frozen in liquid nitrogen and grounded to fine powder. The total pigments were extracted in 80% acetone buffered with Tris-HCl pH 7.4 and the chlorophyll a and b content measured by multi-wave length absorbance according to (Porra, 2002).

### Protein analysis and immunodetection

Protein samples were prepared from entire 14 days-old plantlets. Each sample was composed of at least three individuals, the fresh weight recorded and the plantlets were then fresh frozen in liquid nitrogen. The frozen samples were ground to fine powder using glass beads mixing in a Ivoclar Vivadent shaker (Silamat) two times for 10 seconds. The extraction was performed by homogenizing the sample powder in lysis buffer containing 100 mM Tris-HCl, pH 7.8, 2% SDS, 50 mM NaF, and 1× Protease Inhibitor Cocktail for Plant (Sigma-Aldrich) and then incubating for 30 min at 37°C. The supernatant was clarified by centrifugation and add to a mix composed by 25% of the proteins sample, 50% of deionized water and 25% of 4x sample buffer (0.2 M Tris/HCl pH 6.8, 0.4 M Dithiotreitol, 8% w/v SDS, 0.4 % w/v Bromophenol Blue, 40 % v/v Glycerol), which was then used loaded on a Tris-Gly SDS-PAGE 12% acrylamide gel. For the separation of the phosphorylated form of the protein, we used Tris-Gly gels containing 30 μM Phos-tag (Wako Chemicals) and 60 μM MnCl2. The proteins were separated by electrophoresis on the gel and transferred to a nitrocellulose membrane for immuno-detection. The nitrocellulose membranes were blocked with commercial skim-milk 5% (M) in TBS Tween (0.25%), except for the membranes used for the detection of LHCB2, which were blocked with 3% BSA (Applichem) in TBS Triton X-100 (0.1%) to allow the membrane dephosphorylation, detailed in (P. Longoni et al., 2015). After the blocking the membranes were decorated with primary antibodies for the detection of relevant proteins. The antibodies recognizing the following proteins were obtained from Agrisera: LHCB1(AS09 522), LHCB2 (AS01 003), LHCB2-P (AS13 2705), LHCB3 (AS01 002), LHCB4 (CP29)(AS04 045), LHCB5 (CP26) (AS01 009), LHCB6 (CP24)(AS04 010), D1(PsbA) (AS05 084), PsbA-P (AS13 2669), D2 (PsbD) (AS06 146), PsbC (CP43)(AS06 111), PsbB (CP47) (AS04 038), PETC (AS08 330), PsaB (AS10 695), PsaC (AS10 939), PSAD (AS09 461), PSAH (AS06 105), LHCA1 (AS01 005), LHCA2 (AS01 006), LHCA3 (AS01 007), LHCA4 (AS01 008), STN7 (AS16 4098), PPH1 (AS16 4084), STN8 (AS10 1601), ATPC (Agrisera, AS08 312). The antibody recognizing ACT2 (Actin) (A0480) was from Sigma-Aldrich, finally the anti PBCP antibody was a gift from Michel Goldschmidt-Clermont. Secondary antibodies (anti-rabbit (Merck, AP132P) or anti-mouse (Sigma, A5278) conjugated with HRP were used for the detection of the primary antibody by enhanced chemiluminescence (ECL) the light signal was recorded using an imager for chemiluminescence (Amersham Imager 600, Amersham Biosciences, Inc). Band intensity was measured with ImageQuant software (Amersham).

### Thylakoid preparation and fractionation

Thylakoid preparation was performed according to (Arnold et al., 2014), from adult (four weeks old) full rosettes. Separation of supercomplexes by blue native polyacrylamide gel electrophoresis was performed as previously described (Järvi, Suorsa, Paakkarinen, & Aro, 2011), using SERVAGel™ N 3 - 12, Vertical Native Gels (Serva). The grana stacks and the stroma lamellae fraction were obtained by solubilization of the thylakoid preparation (0.5 mg mL^−1^) with 1% water soluble digitonin (Serva), followed by sequential ultracentrifugation 40’000xg for 40 min to pellet the grana and 180’000xg for 90 min to pellet the stroma lamellae fraction.

### Lipid analysis

The total lipid extraction was performed on samples of three seedlings. After measurement of the total fresh weight (FW), the samples were flash frozen in liquid nitrogen and stored at −80°C. The frozen samples were ground to fine powder using glass beads mixing in an Ivoclar Vivadent shaker (Silamat) two times for 10 seconds. Total lipids were extracted from the powder by adding 10 ul of a tetrahydrofuran: methanol 50:50 (v/v) solution per mg of FW. Plants debris were pelleted by centrifugation (3 min, 14,000 × g); finally, clear supernatant was pipetted to an HPLC vial to perform an ultra-high pressure liquid chromatography separation followed by atmospheric pressure chemical ionization-quadrupole time-of-flight mass spectrometry (UHPLC-APCI-QTOF-MS) according to (Sattari Vayghan et al., 2020; Spicher, Glauser, & Kessler, 2016). Briefly, the separation was performed on a reverse-phase Acquity BEH C18 column (50 × 2.1 mm, 1.7 μm) maintained at 60°C. The elution profile was: solvent A = water; solvent B = methanol; 80–100% B in 3 min, 100% B for 2 min, re-equilibration at 80% B for 2.0 min, the flow rate 0.8 ml min–1 and the injection volume 2.5 μl. Mass data were acquired using MassLynx version 4.1 (Waters) and TargetLynx (Waters) was used for processing. Compound identity was determined based on reference standards which were used for the quantification curves as well (Spicher et al., 2016). Lutein and zeaxanthin as well as violaxanthin and neoxanthin could be resolved neither by chromatography nor by mass spectrometry under the conditions employed; therefore, the measured peak corresponds to the sum of both compounds.

### Chlorophyll fluorescence analysis

Before the measurements, the plants were dark acclimated for at least 15 minutes. The chlorophyll fluorescence was recorded with a MF800 Fluorcam (Photon System Instrument, Czechia) using a personalized light protocol (Pralon et al., 2020). Briefly, the maximum fluorescence of dark adapted plants was first measured during a saturating light pulse (F_M_). Subsequently, the plants were exposed to steps of increasing blue light intensity. The length of the steps was 1 minute for the seedling screening (92, 208, 335, 846, 1365, 1878 μmol photons m^−2^ s^−1^ of PAR intensity), and 5 minutes for the analyses on adult plants (40, 95, 150, 380, 620, 850 μmol photons m^−2^ s^−1^ of PAR intensity). At the end of each step the maximal fluorescence of light acclimated plants (F_M_’) was measured during a saturating light pulse. After each step the actinic light was turned off, the plants were then exposed for 2s to far-red light to oxidize the photosynthetic ETC, after that the recorder fluorescence value was used as F_0_’. To calculate the derived parameters we used the fluorescence recorded just before the saturating light pulse as F_S_. The non-photochemical energy dissipation was measured as NPQ = (F_M_–F_M_’)/F_M_’. PSII quantum yield under light (ΦPSII) was calculated as ΦPSII = (F_M_’–F_S_)/F_M_’. The fraction of the closed PSII centers (1-qL) was calculated with the following formula: 1-qL = 1-[(F_M_’–F_S_)/(F_M_’–F_0_’)]* (F_0_’/F_S_) (Kramer et al., 2004). The chlorophyll fluorescence was also measured with a portable device (Multispeq v2, PhotosynQ, Firmware 2.1), with a modified protocol available at this address https://photosynq.org/protocols/npq_phi2_from_simple_fluor_light_curve_with_recovery. Due to the LED limitations of this instrument the time for each light intensity was reduced to 30 seconds (40, 95, 150, 380, 620, 850 μmol photons m^−2^ s^−1^ of PAR intensity).

Rapid chlorophyll fluorescence induction was measured on detached leaves with the Plant Efficiency Analyser (M-PEA 2; Hansatech Ltd). The following protocol was used: after an initial saturating pulse (3000 μmol photons m^−2^ s^−1^, 700 ms, red light, dominant λ625 nm), the same pulse was repeated after sequentially longer dark intervals (0.05, 4, 8, 12, 16, 20, and 24 s) for a total of eight pulses. After the eighth pulse, far-red light (20%) was turned on, and the sample was submitted to a second series of saturating pulses separated by increasing far-red light intervals (0.05, 0.1, 0.2, 0.4, 0.8, and 1.6 s). For each pulse, the fast chlorophyll fluorescence curve was extrapolated by the M-PEA 2 software (Hansatech Ltd). The variable fluorescence at 3 ms (Vj) was analyzed by plotting as a function of the time between pulses.

To assess the functional antenna size of PSII, fluorescence induction measurements upon a transition from darkness to low light were performed on DCMU (3-(3,4-dichlorophenyl)-1,1-dimethylurea)-infiltrated leaves according to (Nicol et al., 2019). In brief, in the presence of DCMU, the rise to the fluorescence maximum (F_M_) represents a single, stable charge separation in PSII, the rate of the latter being directly proportional to the PSII absorption cross-section at a given light intensity. The reciprocal of the integrated area above the induction curve yields the maximal initial rate of PSII (when all PSII are open).

FLIM measurements were performed on whole leaves incubated with DCMU with the same setup as the confocal images (see Confocal Microscopy), except that it was coupled to a PicoHarp 300 time-correlated single photon counting (TCSPC) module (PicoQuant, Berlin, Germany) as described in (Wientjes et al., 2017). Sample were excited with 633 nm light (40 MHz rep. rate) and fluorescence emission was detected at 710-750 nm. Each image was recorded for 1 min. The total image size was 116μm*116μm, with 128*128 pixels. The time step for the TCSPC detection was 32 ps/channel. Decay traces were fitted with three exponentials F(t)= a1* e^(-t/τ1) + a2* e^(-t/τ2) + a3* e^(-t/τ3) with (a) the amplitude and (τ) the lifetimes (τ1=100 ps, τ2=900 ps and τ3=2 ns) using the FLIMfit software tool (Warren et al., 2013). Based on the amplitude of the PSI lifetime (100ps), the ratio of chlorophylls associated with PSI compared to PSII can be calculated according to PSI/(PSI+PSII)=c*a1/(1-a1+c*a1) with c being a previously determined correction factor for this specific setup (c=0.49) (Wientjes et al., 2017).

### Absorption spectroscopy in vivo

For the calculation of the ratio of CEF over LEF we used the protocol described in Kramer et al. 2020. Briefly, the plants were grown under long day conditions and initially dark incubated for 60 min and then 5min before transfer into actinic light with 630 μmol photons m^−2^ s^−1^ PAR for light acclimation. After 10, 20, 40, 60 s or 5 min of illumination, LEF and CEF were measured by following the relaxation kinetics of the carotenoid electrochromic bandshift at 520 nm (corrected for the signal at 546 nm) using a JTS-10 spectrophotometer (Biologic, France). The ECS spectral change is a shift in the pigment absorption bands that is linearly correlated with the light-induced generation of a membrane potential across the thylakoid membranes (Bailleul et al., 2010). Under steady state continuous illumination, the ECS signal stems from transmembrane potential generation by PSII, the cytb6f complex, and PSI and from transmembrane potential dissipation by the ATP synthase CF_0_-F_1_. When light is switched off, reaction centres’ activity stops immediately, while ATPase and the cytb6f complex activities remain (transiently) unchanged. Therefore, the initial rate of ECS decay is proportional to the rate of PSI and PSII photochemistry (*i.e*. to the rate of ‘total’ electron flow). This can be calculated by dividing this rate (expressed as -ΔI/I per unit of time) by the amplitude of the ECS signal (again expressed as -ΔI/I) induced by the transfer of one charge across the membrane (e.g. one PSI turnover). The rate of CEF can be evaluated using the same approach under conditions where PSI only is excited by exposure to saturating far red light (λ >720 nm) for enough time to be at the steady-state (5min in our condition). Results were expressed as electrons^−1^ s^−1^ and estimated from the amplitude of the electrochromic shift signal upon excitation with a saturating single turnover flash (5 ns Nd:YAG laser flash, Continuum Minilite, 532 nm flash exciting 4-(dicyanomethylene)-2-methyl-6-(p-dimethylaminostyryl)-4H-pyran (DCM), emission peak at 630 nm). Total electron flow was measured following a pulse of actinic light (λ = 640 ± 20 nm FWHM) at 1100 μmol photons m^−2^ s^−1^ while CEF was measured with a pulse of far red light at the maximum setting (estimated as 1400 μmol photons m^−2^ s^−1^ by the manufacturer).

Functional antenna size of PSII and PSI was assessed using ECS as described in (Hu, Nawrocki, & Croce, 2021). In brief, similarly to the light-to-dark transition described above, the initial slope of ECS signal during onset of light is directly proportional to the absorption-limited, maximal rate of charge separation by fully open PSII+PSI. Addition of DCMU and hydroxylamine by vacuum infiltration of the leaf inhibited PSII activity (systematically verified with fluorescence measurements), allowing to separately quantify the rates of both PSII and PSI. The rates in e^−^ s^−1^ PSI^−1^ (or e^−^ s^−1^ PSII^−1^) were obtained by dividing the slope by the ECS signal obtained 140 μs after a saturating, single-turnover laser flash (see above) corresponding to 1 charge separation PS^−1^. Note that the absolute value of maximal PSII rate at 80 μmol photons m^−2^ s^−1^ obtained with both fluorescence- and ECS-based methods is identical (Fig. 4A).

### Confocal microscopy

Confocal images were recorded on an (inverted) confocal Leica TCS SP8 system equipped with a 63 × 1.20 NA water immersion objective. Chloroplasts were freshly isolated and chlorophylls were excited at 633 nm with a pulsed laser (40 MHz) and an intensity of 100 nW. The fluorescence was detected at 650-680nm (PSII-maximum) and 710-750nm (PSI-maximum) with internal hybrid detectors. The total image size was 9.2 μm * 9.2 μm, with 128*128 pixels.

### Transmission electron microscopy

Samples preparation and analysis were performed as previously described (Martinis, Glauser, Valimareanu, & Kessler, 2013), with minor modifications. Leaves from 14 days-old WT and L1ko plants were fixed in fixative buffer (5% (W/V) glutaraldehyde and 4% (W/V) formaldehyde in 100 mM phosphate buffer (pH 6.8)) overnight at 4°C, rinsed several times in phosphate buffer, and post fixed for 2 h with 1% (W/V) osmium tetroxide in phosphate buffer at 20°C. After two further washing steps in phosphate buffer and distilled water, the samples were dehydrated in ethanol and embedded in Spurr’s low-viscosity resin (Polyscience). Ultrathin sections of 50–70 nm were cut with a diamond knife (Ultracut-E microtome-Reihtert-Jung), mounted on uncoated copper grids. The sections were post stained with Uranyless and Reynold’s lead citrate stain (Delta Microscopies). Sections were observed with a Philips CM 100 transmission electron microscope operating at 60 kV (Philips Electron Optics BV, Eindhoven, the Netherlands).

### Statistical analysis

The normal distribution of the residuals of each data set was tested before any other statistical analysis. If this assumption was met, an ANOVA model was utilized; otherwise, a Kruskal– Wallis rank sum test was performed. If the results were significant, we used post hoc Student’s T-test for multiple comparisons. The reported p-values were obtained with the latter. The effect of genotypes on the correlation between the number of stacks per grana or grana width was tested using analyses of covariance, with number of stacks as response variable, and the genotypes by grana, or by grana width as factors. We used a generalised linear model (GLM) with poisson distribution for assessing the numbers of stacks, and a linear model for grana with. The calculations were performed with RStudio (Version 1.2.5019 RStudio Inc).

## Data availability

The raw data and the pictures used for this manuscript are available at https://zenodo.org/record/5729177#.YaDZILrjKUk. Further data, and the described lines, can be provided by the authors upon reasonable request.

## Author contributions

H.S.V and F.L designed the experimental plan. R.C, W.N and C.H measured the photochemical rate of photosystems. G.F. and D.T. performed the CEF/LEF analysis. H.S.V and V.D. produced the electron microscopy pictures. E.W. analyzed the images. E.W and C.S performed confocal microscopy and FLIM analysis. H.S.V and F.L performed the other experiments. G.G performed the lipid profile analysis. F.L. performed the statistical analysis of the data. All authors contributed in the redaction. All authors read and approved the manuscript.

## Acknowledgments

H.S.V. and F.L. work was supported by the University of Neuchâtel and the Swiss National Science Foundation (31003A_179417).

The authors thanks prof. Sergio Rasmann for his assistance in the statistical analysis of the grana distribution, and Jenny Pego Magalhaes for her technical help in TEM analysis.

---

**Table 1.**
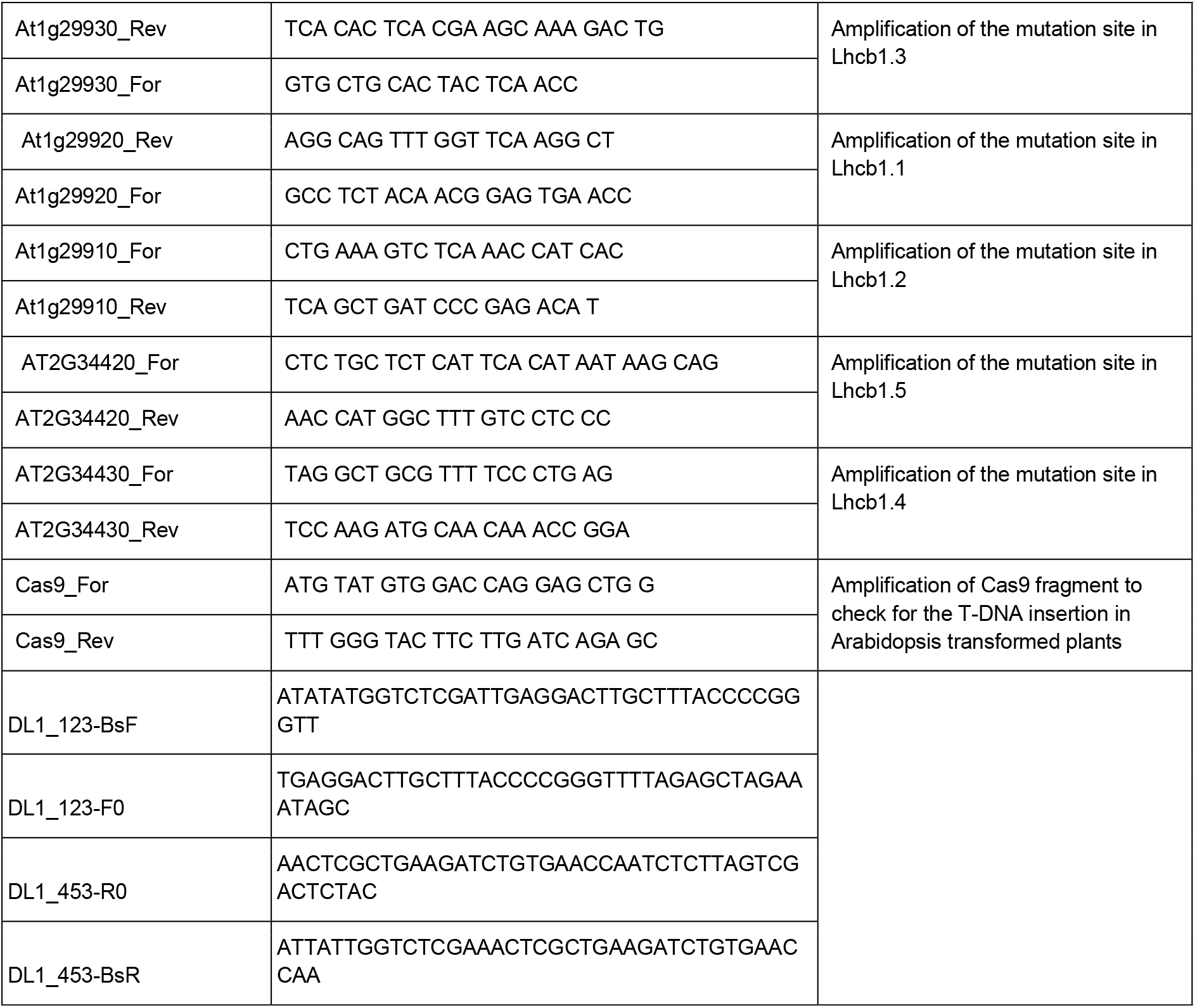
oligonucleotides used in this report.

## Notes

https://zenodo.org/record/5729177#.YaDZILrjKUk

### Competing Interest Statement

The authors have declared no competing interest.

## References

Albanese, P., Tamara, S., Saracco, G., Scheltema, R. A., & Pagliano, C. (2020). How paired PSII-LHCII supercomplexes mediate the stacking of plant thylakoid membranes unveiled by structural mass-spectrometry. Nat Commun, 11(1), 1361. doi:10.1038/s41467-020-15184-1

Allen, J. F. (1992). Protein phosphorylation in regulation of photosynthesis. Biochimica et Biophysica Acta (BBA)-Bioenergetics, 1098(3), 275–335.

Ancin, M., Fernandez-San Millan, A., Larraya, L., Morales, F., Veramendi, J., Aranjuelo, I., & Farran, I. (2019). Overexpression of thioredoxin m in tobacco chloroplasts inhibits the protein kinase STN7 and alters photosynthetic performance. J Exp Bot, 70(3), 1005–1016. doi:10.1093/jxb/ery415

Andersson, J., Wentworth, M., Walters, R. G., Howard, C. A., Ruban, A. V., Horton, P., & Jansson, S. (2003). Absence of the Lhcb1 and Lhcb2 proteins of the light-harvesting complex of photosystem II - effects on photosynthesis, grana stacking and fitness. Plant J, 35(3), 350–361. doi:10.1046/j.1365-313x.2003.01811.x

Armbruster, U., Labs, M., Pribil, M., Viola, S., Xu, W., Scharfenberg, M., … Rappaport, F. (2013). Arabidopsis CURVATURE THYLAKOID1 proteins modify thylakoid architecture by inducing membrane curvature. The Plant Cell, 25(7), 2661–2678.

Arnold, J., Shapiguzov, A., Fucile, G., Rochaix, J.-D., Goldschmidt-Clermont, M., & Eichacker, L. A. (2014). Separation of membrane protein complexes by native LDS-PAGE. In Plant Proteomics (pp. 667–676): Springer.

Ballottari, M., Girardon, J., Dall’Osto, L., & Bassi, R. (2012). Evolution and functional properties of photosystem II light harvesting complexes in eukaryotes. Biochimica et Biophysica Acta (BBA)-Bioenergetics, 1817(1), 143–157.

Barber, J. (1980). An explanation for the relationship between salt-induced thylakoid stacking and the chlorophyll fluorescence changes associated with changes in spillover of energy from photosystem II to photosystem I. FEBS letters, 118(1), 1–10.

Bellafiore, S., Barneche, F., Peltier, G., & Rochaix, J. D. (2005). State transitions and light adaptation require chloroplast thylakoid protein kinase STN7. Nature, 433(7028), 892–895. doi:10.1038/nature03286

Bos, I., Bland, K. M., Tian, L., Croce, R., Frankel, L. K., van Amerongen, H., … Wientjes, E. (2017). Multiple LHCII antennae can transfer energy efficiently to a single Photosystem I. Biochimica et Biophysica Acta (BBA)-Bioenergetics, 1858(5), 371–378.

Bos, P., Oosterwijk, A., Koehorst, R., Bader, A., Philippi, J., van Amerongen, H., & Wientjes, E. (2019). Digitonin-sensitive LHCII enlarges the antenna of Photosystem I in stroma lamellae of Arabidopsis thaliana after far-red and blue-light treatment. Biochimica et Biophysica Acta (BBA)-Bioenergetics, 1860(8), 651–658.

Caffarri, S., Croce, R., Cattivelli, L., & Bassi, R. (2004). A look within LHCII: differential analysis of the Lhcb1-3 complexes building the major trimeric antenna complex of higher-plant photosynthesis. Biochemistry, 43(29), 9467–9476. doi:10.1021/bi036265i

Chukhutsina, V. U., Liu, X., Xu, P., & Croce, R. (2020). Light-harvesting complex II is an antenna of photosystem I in dark-adapted plants. Nature Plants, 6(7), 860–868. doi:10.1038/s41477-020-0693-4

Crepin, A., & Caffarri, S. (2015). The specific localizations of phosphorylated Lhcb1 and Lhcb2 isoforms reveal the role of Lhcb2 in the formation of the PSI-LHCII supercomplex in Arabidopsis during state transitions. Biochim Biophys Acta, 1847(12), 1539–1548. doi:10.1016/j.bbabio.2015.09.005

Crepin, A., & Caffarri, S. (2018). Functions and evolution of Lhcb isoforms composing LHCII, the major light harvesting complex of photosystem II of green eukaryotic organisms. Current Protein and Peptide Science, 19(7), 699–713.

Crepin, A., Kucerova, Z., Kosta, A., Durand, E., & Caffarri, S. (2020). Isolation and characterization of a large photosystem I-light-harvesting complex II supercomplex with an additional Lhca1-a4 dimer in Arabidopsis. Plant J, 102(2), 398–409. doi:10.1111/tpj.14634

Croce, R., & Amerongen, H. v. (2020). Light harvesting in oxygenic photosynthesis: Structural biology meets spectroscopy. Science, 369(6506), eaay2058. doi:doi:10.1126/science.aay2058

Daum, B., Nicastro, D., Austin, J., McIntosh, J. R., & Kühlbrandt, W. (2010). Arrangement of photosystem II and ATP synthase in chloroplast membranes of spinach and pea. The Plant Cell, 22(4), 1299–1312.

Dekker, J. P., & Boekema, E. J. (2005). Supramolecular organization of thylakoid membrane proteins in green plants. Biochimica et Biophysica Acta (BBA)-Bioenergetics, 1706(1-2), 12–39.

Durr, J., Papareddy, R., Nakajima, K., & Gutierrez-Marcos, J. (2018). Highly efficient heritable targeted deletions of gene clusters and non-coding regulatory regions in Arabidopsis using CRISPR/Cas9. Scientific reports, 8(1), 1–11.

Fristedt, R., Willig, A., Granath, P., Crevecoeur, M., Rochaix, J.-D., & Vener, A. V. (2009). Phosphorylation of photosystem II controls functional macroscopic folding of photosynthetic membranes in Arabidopsis. The Plant Cell, 21(12), 3950–3964.

Galka, P., Santabarbara, S., Khuong, T. T. H., Degand, H., Morsomme, P., Jennings, R. C., … Caffarri, S. (2012). Functional analyses of the plant photosystem I–light-harvesting complex II supercomplex reveal that light-harvesting complex II loosely bound to photosystem II is a very efficient antenna for photosystem I in state II. The Plant Cell, 24(7), 2963–2978.

Goral, T. K., Johnson, M. P., Duffy, C. D., Brain, A. P., Ruban, A. V., & Mullineaux, C. W. (2012). Light-harvesting antenna composition controls the macrostructure and dynamics of thylakoid membranes in Arabidopsis. The Plant Journal, 69(2), 289–301.

Heigwer, F., Kerr, G., & Boutros, M. (2014). E-CRISP: fast CRISPR target site identification. Nature methods, 11(2), 122–123.

Hepworth, C., Wood, W. H. J., Emrich-Mills, T. Z., Proctor, M. S., Casson, S., & Johnson, M. P. (2021). Dynamic thylakoid stacking and state transitions work synergistically to avoid acceptor-side limitation of photosystem I. Nat Plants, 7(1), 87–98. doi:10.1038/s41477-020-00828-3

Hobe, S., Foerster, R., Klingler, J., & Paulsen, H. (1995). N-proximal sequence motif in light-harvesting chlorophyll a/b-binding protein is essential for the trimerization of light-harvesting chlorophyll a/b complex. Biochemistry, 34(32), 10224–10228.

Horton, P., & Ruban, A. (2005). Molecular design of the photosystem II light-harvesting antenna: photosynthesis and photoprotection. J Exp Bot, 56(411), 365–373. doi:10.1093/jxb/eri023

Hu, C., Nawrocki, W. J., & Croce, R. (2021). Long-term adaptation of Arabidopsis thaliana to far-red light. Plant Cell Environ, 44(9), 3002–3014. doi:10.1111/pce.14032

Jansson, S. (1999). A guide to the Lhc genes and their relatives in Arabidopsis. Trends in plant science, 4(6), 236–240.

Jansson, S., Pichersky, E., Bassi, R., Green, B. R., Ikeuchi, M., Melis, A., … Thornber, J. P. (1992). A nomenclature for the genes encoding the chlorophyll a/b-binding proteins of higher plants. Plant Molecular Biology Reporter, 10(3), 242–253.

Järvi, S., Suorsa, M., Paakkarinen, V., & Aro, E.-M. (2011). Optimized native gel systems for separation of thylakoid protein complexes: novel super-and mega-complexes. Biochemical Journal, 439(2), 207–214.

Johnson, M. P., Vasilev, C., Olsen, J. D., & Hunter, C. N. (2014). Nanodomains of Cytochrome b 6 f and Photosystem II Complexes in Spinach Grana Thylakoid Membranes The Plant Cell, 26(7), 3051–3061. doi:10.1105/tpc.114.127233

Johnson, M. P., & Wientjes, E. (2020). The relevance of dynamic thylakoid organisation to photosynthetic regulation. Biochim Biophys Acta Bioenerg, 1861(4), 148039. doi:10.1016/j.bbabio.2019.06.011

Kirchhoff, H., Mukherjee, U., & Galla, H.-J. (2002). Molecular architecture of the thylakoid membrane: lipid diffusion space for plastoquinone. Biochemistry, 41(15), 4872–4882.

Klimmek, F., Sjodin, A., Noutsos, C., Leister, D., & Jansson, S. (2006). Abundantly and rarely expressed Lhc protein genes exhibit distinct regulation patterns in plants. Plant Physiol, 140(3), 793–804. doi:10.1104/pp.105.073304

Kouřil, R., Zygadlo, A., Arteni, A. A., de Wit, C. D., Dekker, J. P., Jensen, P. E., … Boekema, E. J. (2005). Structural characterization of a complex of photosystem I and light-harvesting complex II of Arabidopsis thaliana. Biochemistry, 44(33), 10935–10940.

Kramer, D. M., Johnson, G., Kiirats, O., & Edwards, G. E. (2004). New Fluorescence Parameters for the Determination of QA Redox State and Excitation Energy Fluxes. Photosynth Res, 79(2), 209. doi:10.1023/B:PRES.0000015391.99477.0d

Krause, G. H., & Weis, E. (1984). Chlorophyll fluorescence as a tool in plant physiology. Photosynthesis Research, 5(2), 139–157. doi:10.1007/BF00028527

Leoni, C., Pietrzykowska, M., Kiss, A. Z., Suorsa, M., Ceci, L. R., Aro, E. M., & Jansson, S. (2013). Very rapid phosphorylation kinetics suggest a unique role for Lhcb2 during state transitions in Arabidopsis. Plant J, 76(2), 236–246. doi:10.1111/tpj.12297

Leutwiler, L. S., Meyerowitz, E. M., & Tobin, E. M. (1986). Structure and expression of three light-harvesting chlorophyll a/b-binding protein genes in Arabidopsis thaliana. Nucleic Acids Res, 14(10), 4051–4064. doi:10.1093/nar/14.10.4051

Li, Z., Ahn, T. K., Avenson, T. J., Ballottari, M., Cruz, J. A., Kramer, D. M., … Niyogi, K. K. (2009). Lutein Accumulation in the Absence of Zeaxanthin Restores Nonphotochemical Quenching in the Arabidopsis thaliana npq1 Mutant The Plant Cell, 21(6), 1798–1812. doi:10.1105/tpc.109.066571

Longoni, F. P., & Goldschmidt-Clermont, M. (2021). Thylakoid Protein Phosphorylation in Chloroplasts. Plant Cell Physiol. doi:10.1093/pcp/pcab043

Longoni, P., Douchi, D., Cariti, F., Fucile, G., & Goldschmidt-Clermont, M. (2015). Phosphorylation of the Light-Harvesting Complex II Isoform Lhcb2 Is Central to State Transitions. Plant Physiol, 169(4), 2874–2883. doi:10.1104/pp.15.01498

Longoni, P., Samol, I., & Goldschmidt-Clermont, M. (2019). The Kinase STATE TRANSITION 8 Phosphorylates Light Harvesting Complex II and Contributes to Light Acclimation in Arabidopsis thaliana. Front Plant Sci, 10, 1156. doi:10.3389/fpls.2019.01156

Martinis, J., Glauser, G., Valimareanu, S., & Kessler, F. (2013). A chloroplast ABC1-like kinase regulates vitamin E metabolism in Arabidopsis. Plant Physiology, 162(2), 652–662.

Mazor, Y., Borovikova, A., Caspy, I., & Nelson, N. (2017). Structure of the plant photosystem I supercomplex at 2.6 Å resolution. Nature Plants, 3(3), 17014. doi:10.1038/nplants.2017.14

McGrath, J. M., Terzaghi, W. B., Sridhar, P., Cashmore, A. R., & Pichersky, E. (1992). Sequence of the fourth and fifth Photosystem II type I chlorophyll a/b-binding protein genes of Arabidopsis thaliana and evidence for the presence of a full complement of the extended CAB gene family. Plant Mol Biol, 19(5), 725–733. doi:10.1007/BF00027069

Montague, T. G., Cruz, J. M., Gagnon, J. A., Church, G. M., & Valen, E. (2014). CHOPCHOP: a CRISPR/Cas9 and TALEN web tool for genome editing. Nucleic acids research, 42(W1), W401–W407.

Nawrocki, W. J., Tourasse, N. J., Taly, A., Rappaport, F., & Wollman, F.-A. (2015). The plastid terminal oxidase: its elusive function points to multiple contributions to plastid physiology. Annual review of plant biology, 66, 49–74.

Nicol, L., Nawrocki, W. J., & Croce, R. (2019). Disentangling the sites of non-photochemical quenching in vascular plants. Nat Plants, 5(11), 1177–1183. doi:10.1038/s41477-019-0526-5

Ordon, J., Bressan, M., Kretschmer, C., Dall’Osto, L., Marillonnet, S., Bassi, R., & Stuttmann, J. (2020). Optimized Cas9 expression systems for highly efficient Arabidopsis genome editing facilitate isolation of complex alleles in a single generation. Funct Integr Genomics, 20(1), 151–162. doi:10.1007/s10142-019-00665-4

Pan, X., Cao, P., Su, X., Liu, Z., & Li, M. (2020). Structural analysis and comparison of light-harvesting complexes I and II. Biochimica et Biophysica Acta (BBA)-Bioenergetics, 1861(4), 148038.

Pan, X., Ma, J., Su, X., Cao, P., Chang, W., Liu, Z., … Li, M. (2018). Structure of the maize photosystem I supercomplex with light-harvesting complexes I and II. Science, 360(6393), 1109–1113. doi:10.1126/science.aat1156

Peter, G. F., & Thornber, J. P. (1991). Biochemical composition and organization of higher plant photosystem II light-harvesting pigment-proteins. J Biol Chem, 266(25), 16745–16754. Retrieved from https://www.ncbi.nlm.nih.gov/pubmed/1885603

Pietrzykowska, M., Suorsa, M., Semchonok, D. A., Tikkanen, M., Boekema, E. J., Aro, E. M., & Jansson, S. (2014). The light-harvesting chlorophyll a/b binding proteins Lhcb1 and Lhcb2 play complementary roles during state transitions in Arabidopsis. Plant Cell, 26(9), 3646–3660. doi:10.1105/tpc.114.127373

Porra, R. J. (2002). The chequered history of the development and use of simultaneous equations for the accurate determination of chlorophylls a and b. Photosynthesis Research, 73(1), 149–156.

Pralon, T., Collombat, J., Pipitone, R., Ksas, B., Shanmugabalaji, V., Havaux, M., … Kessler, F. (2020). Mutation of the Atypical Kinase ABC1K3 Partially Rescues the PROTON GRADIENT REGULATION 6 Phenotype in Arabidopsis thaliana. Front Plant Sci, 11, 337. doi:10.3389/fpls.2020.00337

Pribil, M., Pesaresi, P., Hertle, A., Barbato, R., & Leister, D. (2010). Role of plastid protein phosphatase TAP38 in LHCII dephosphorylation and thylakoid electron flow. PLoS Biol, 8(1), e1000288. doi:10.1371/journal.pbio.1000288

Qin, X., Suga, M., Kuang, T., & Shen, J.-R. (2015). Structural basis for energy transfer pathways in the plant PSI-LHCI supercomplex. Science, 348(6238), 989–995.

Rantala, M., Rantala, S., & Aro, E. M. (2020). Composition, phosphorylation and dynamic organization of photosynthetic protein complexes in plant thylakoid membrane. Photochem Photobiol Sci, 19(5), 604–619. doi:10.1039/d0pp00025f

Ruban, A. V., Wentworth, M., Yakushevska, A. E., Andersson, J., Lee, P. J., Keegstra, W., … Horton, P. (2003). Plants lacking the main light-harvesting complex retain photosystem II macro-organization. Nature, 421(6923), 648–652. doi:10.1038/nature01344

Sattari Vayghan, H., Tavalaei, S., Grillon, A., Meyer, L., Ballabani, G., Glauser, G., & Longoni, P. (2020). Growth Temperature Influence on Lipids and Photosynthesis in Lepidium sativum. Frontiers in plant science, 11, 745.

Shapiguzov, A., Ingelsson, B., Samol, I., Andres, C., Kessler, F., Rochaix, J. D., … Goldschmidt-Clermont, M. (2010). The PPH1 phosphatase is specifically involved in LHCII dephosphorylation and state transitions in Arabidopsis. Proc Natl Acad Sci U S A, 107(10), 4782–4787. doi:10.1073/pnas.0913810107

Spicher, L., Glauser, G., & Kessler, F. (2016). Lipid antioxidant and galactolipid remodeling under temperature stress in tomato plants. Frontiers in plant science, 7, 167.

Suorsa, M., Rossi, F., Tadini, L., Labs, M., Colombo, M., Jahns, P., … Pesaresi, P. (2016). PGR5-PGRL1-Dependent Cyclic Electron Transport Modulates Linear Electron Transport Rate in Arabidopsis thaliana. Mol Plant, 9(2), 271–288. doi:10.1016/j.molp.2015.12.001

Terao, T., & Katoh, S. (1996). Antenna sizes of photosystem I and photosystem II in chlorophyll b-deficient mutants of rice. Evidence for an antenna function of photosystem II centers that are inactive in electron transport. Plant and Cell Physiology, 37(3), 307–312.

Toth, S. Z., Schansker, G., & Strasser, R. J. (2007). A non-invasive assay of the plastoquinone pool redox state based on the OJIP-transient. Photosynth Res, 93(1-3), 193–203. doi:10.1007/s11120-007-9179-8

Trotta, A., Bajwa, A. A., Mancini, I., Paakkarinen, V., Pribil, M., & Aro, E.-M. (2019). The role of phosphorylation dynamics of CURVATURE THYLAKOID 1B in plant thylakoid membranes. Plant Physiology, 181(4), 1615–1631.

Tyutereva, E. V., Evkaikina, A. I., Ivanova, A. N., & Voitsekhovskaja, O. V. (2017). The absence of chlorophyll b affects lateral mobility of photosynthetic complexes and lipids in grana membranes of Arabidopsis and barley chlorina mutants. Photosynth Res, 133(1-3), 357–370. doi:10.1007/s11120-017-0376-9

Warren, S. C., Margineanu, A., Alibhai, D., Kelly, D. J., Talbot, C., Alexandrov, Y., … French, P. M. (2013). Rapid global fitting of large fluorescence lifetime imaging microscopy datasets. PLoS One, 8(8), e70687. doi:10.1371/journal.pone.0070687

Wientjes, E., & Croce, R. (2011). The light-harvesting complexes of higher-plant Photosystem I: Lhca1/4 and Lhca2/3 form two red-emitting heterodimers. Biochemical Journal, 433(3), 477–485. doi:10.1042/bj20101538

Wientjes, E., Philippi, J., Borst, J. W., & van Amerongen, H. (2017). Imaging the Photosystem I/Photosystem II chlorophyll ratio inside the leaf. Biochimica et Biophysica Acta (BBA)-Bioenergetics, 1858(3), 259–265.

Wientjes, E., van Amerongen, H., & Croce, R. (2013). LHCII is an antenna of both photosystems after long-term acclimation. Biochimica et Biophysica Acta (BBA)-Bioenergetics, 1827(3), 420–426.

Willig, A., Shapiguzov, A., Goldschmidt-Clermont, M., & Rochaix, J. D. (2011). The phosphorylation status of the chloroplast protein kinase STN7 of Arabidopsis affects its turnover. Plant Physiol, 157(4), 2102–2107. doi:10.1104/pp.111.187328

Wunder, T., Xu, W., Liu, Q., Wanner, G., Leister, D., & Pribil, M. (2013). The major thylakoid protein kinases STN7 and STN8 revisited: effects of altered STN8 levels and regulatory specificities of the STN kinases. Front Plant Sci, 4, 417. doi:10.3389/fpls.2013.00417

Xing, H. L., Dong, L., Wang, Z. P., Zhang, H. Y., Han, C. Y., Liu, B., … Chen, Q. J. (2014). A CRISPR/Cas9 toolkit for multiplex genome editing in plants. BMC Plant Biol, 14, 327. doi:10.1186/s12870-014-0327-y

