## Supplementary Figures for "Photosynthetic light harvesting and thylakoid organization in a CRISPR/Cas9 *Arabidopsis thaliana* LHCB1 knockout mutant"

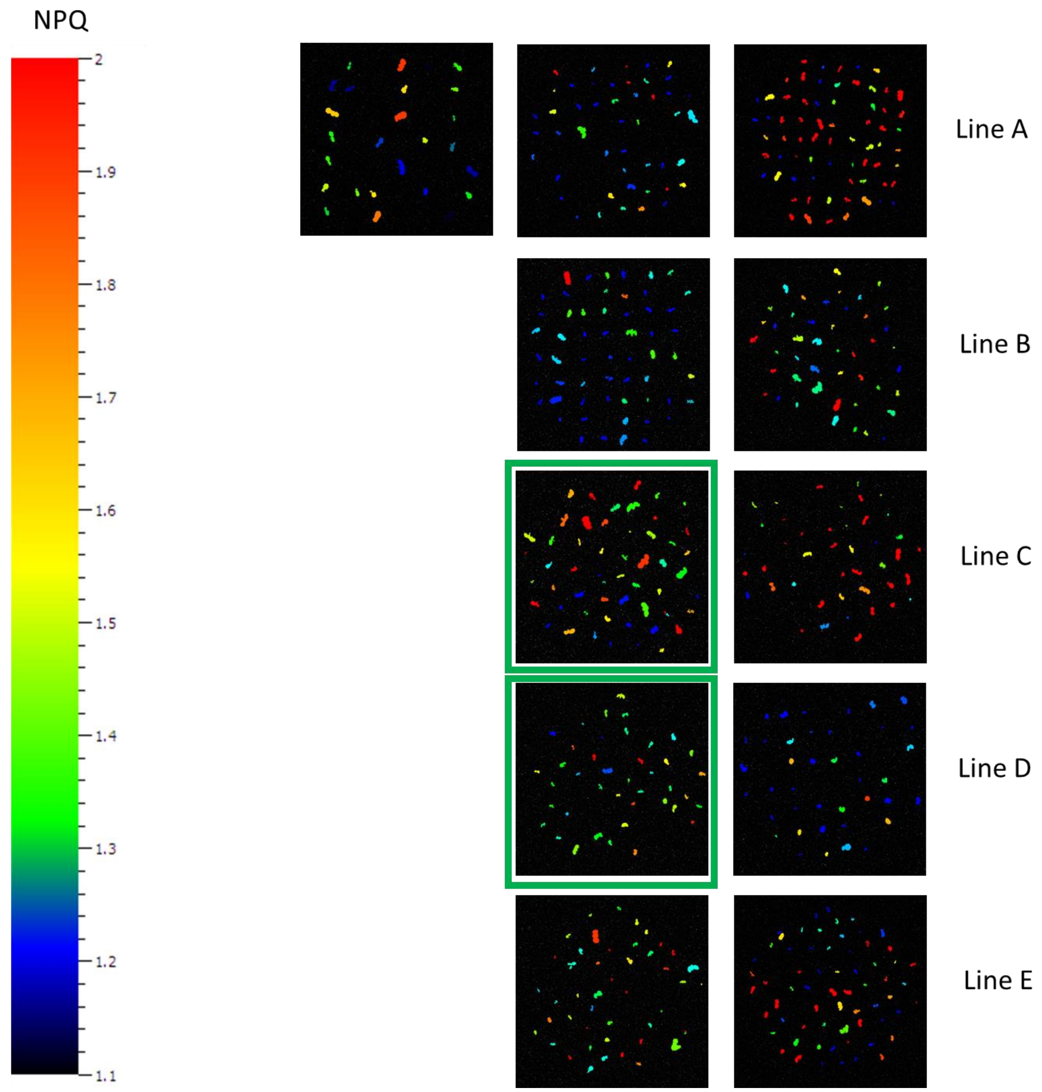


**Figure S1**. Screening of the T2 generation by NPQ. False color image of the calculated NPQ value per plant, the color scale with the corresponding values is shown on the left. Each row shows the plants generated from an independent T0 plant. The range of NPQ values was set between 1.1 (the NPQ value observed in a total Lhcb1 KO mutant) and 2 (the NPQ value of WT plants under this light condition). Plants with a low NPQ were selected from the two sets highlighted with green squares.

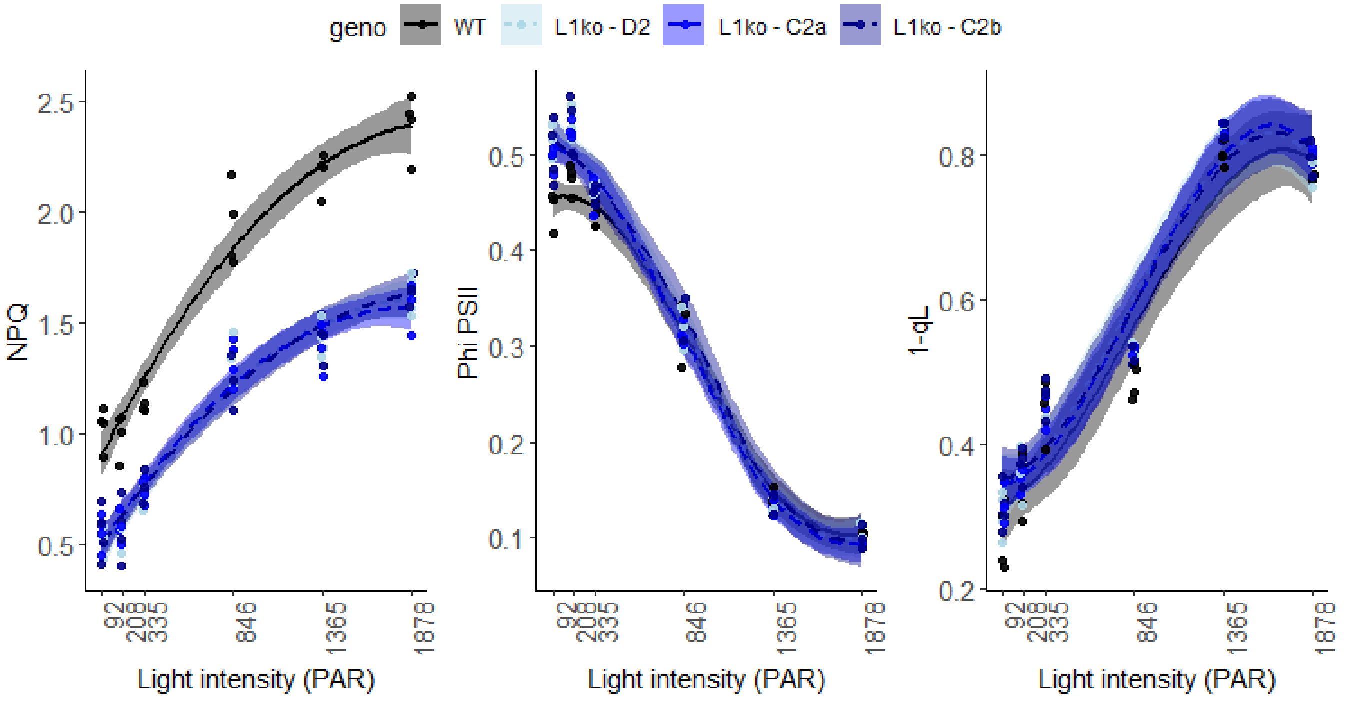


**Figure S2**. Different L1ko mutant lines have the same photosynthetic defects. Room temperature chlorophyll fluorescence analysis. Plates containing 14 days old WT and the three lines of L1ko mutant plantlets were analyzed with increasing actinic light intensities. The non-photochemical quenching (NPQ), the quantum yield of PSII (ΦPSII) and the fraction of closed PSII reaction centers (1-qL), measured at the end of each light step of 1 minute, are shown in the graphs as a function of the light intensity. The WT is shown in black, while the L1ko lines in a gradient of blue. The measured points were fitted with a second degree polynomial function for NPQ, and a third degree polynomial for 1-qL. The points of ΦPSII, were interpolated with a local polynomial regression to visualize a continuous estimate of the standard error. The standard error of the interpolations is shown as a colored area around the curves (n=4).

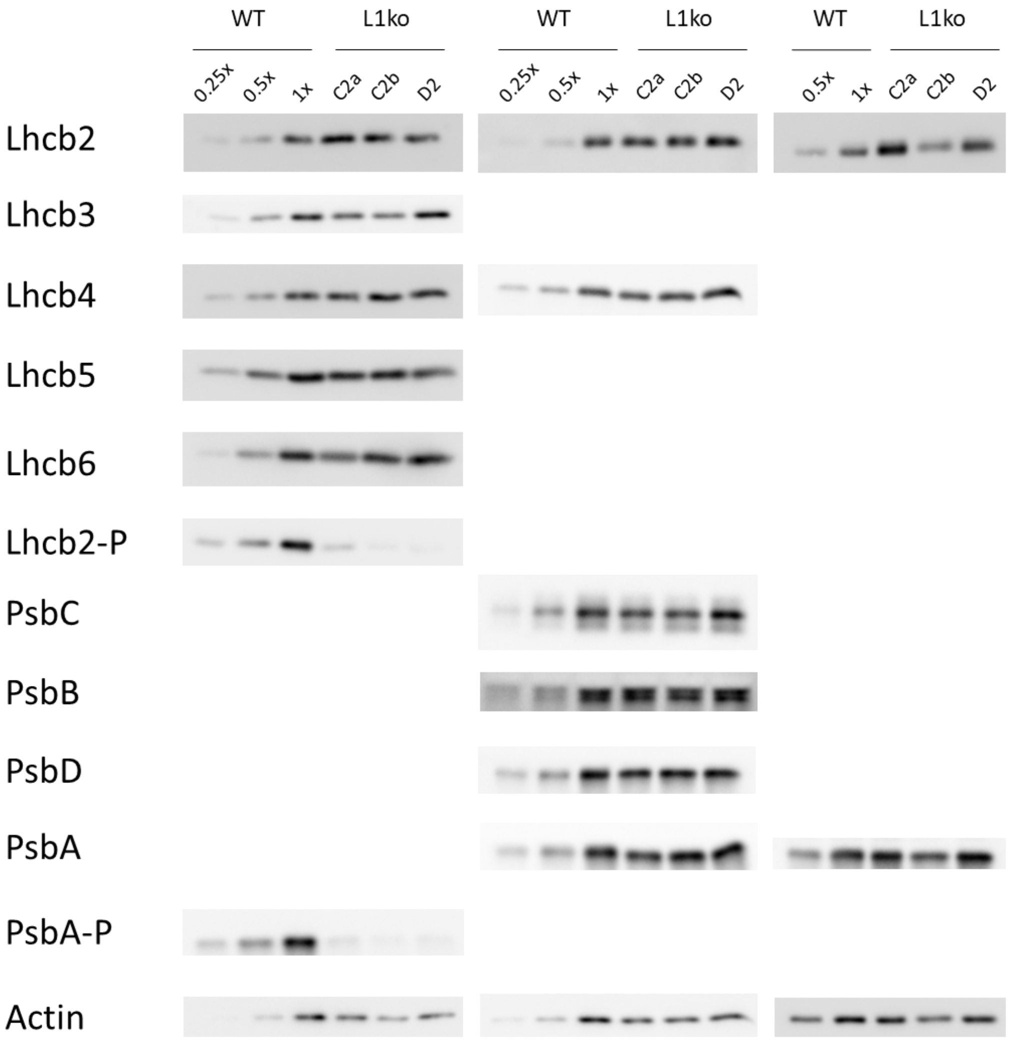


**Figure S3a**. Images of the immunoblot signals analyzed for Figure 3. The relevant protein is indicated in each row, each column represent an independent experiment and set of total protein extracts. WT 0.25x, 0.5x, 1x is the dilution scale used for the quantification of the protein based on signal intensity.

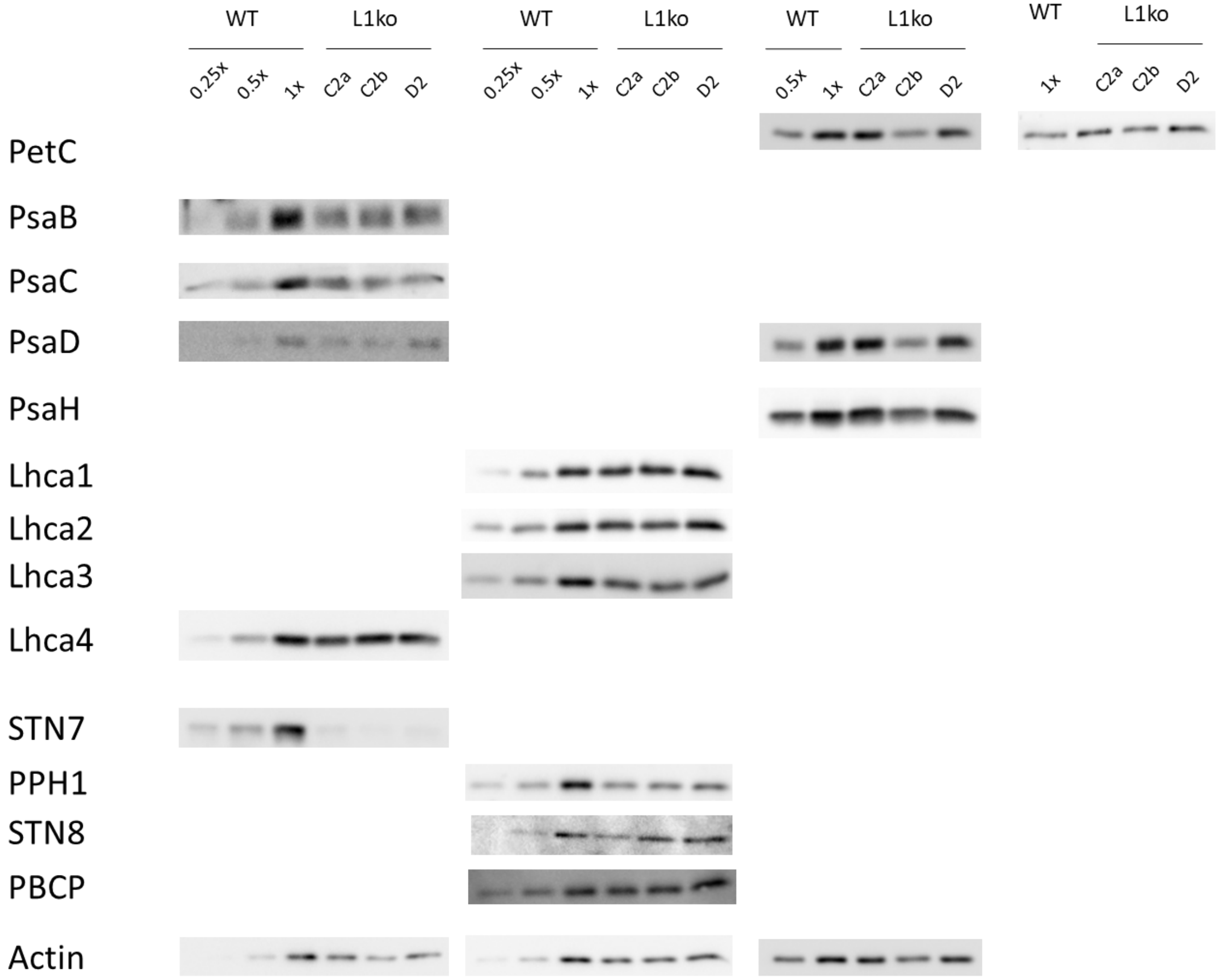


**Figure S3b**. Images of the immunoblot signals analyzed for Figure 3. The relevant protein is indicated in each row, each column represent an independent experiment and set of total protein extracts. WT 0.25x, 0.5x, 1x is the dilution scale used for the quantification of the protein based on signal intensity.

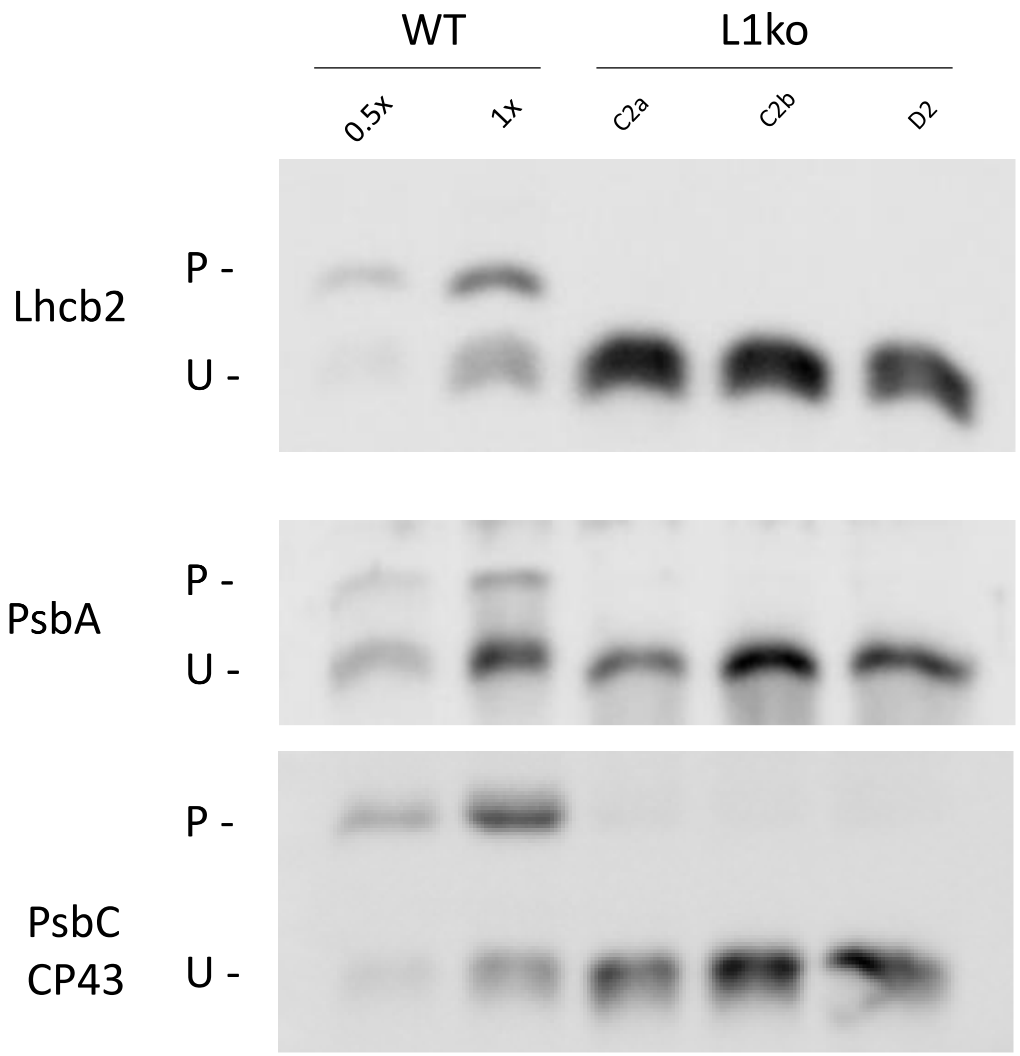


**Figure S4.** Phosphorylation of thylakoid proteins is altered in L1ko lines. Immunoblot of gels separated on a Phos-tag containing gel, separating the phosphorylated form ("P" band, above) from the unphosphorylated protein ("U" band, below). The detected protein is marked on the left, a dilution of the WT was used to assess if the signal was still detectable with 50% of the protein loading (0.5x).

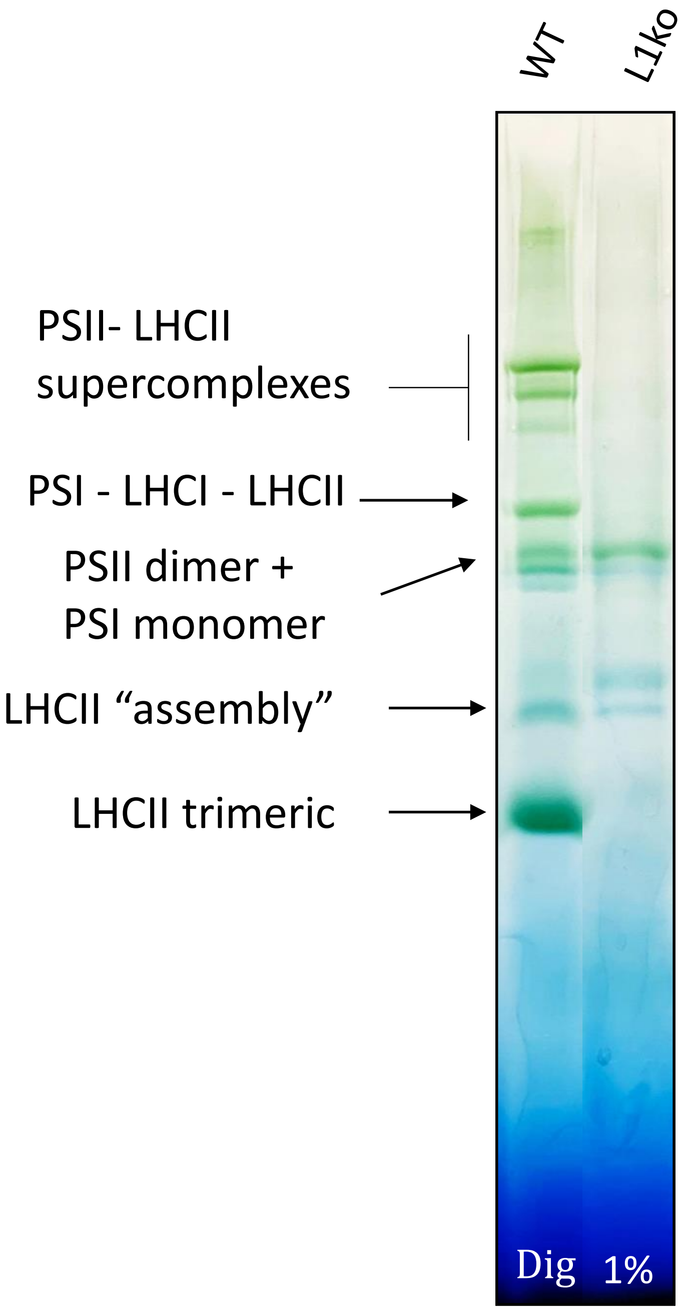


**Figure S5.** Mutation of LHCB1 affects the formation of supercomplexes and the LHCII trimerisation. A preparation of thylakoids from adult leaves of WT and L1ko plants was treated with 1% digitonin to solubilize the membrane and extract the photosynthetic complexes. The separation was performed in a blue native gel, the figure shows a representative lane of the two preparations. The supercomplexes were identified based on previous literature data and their position highlighted with an arrow.

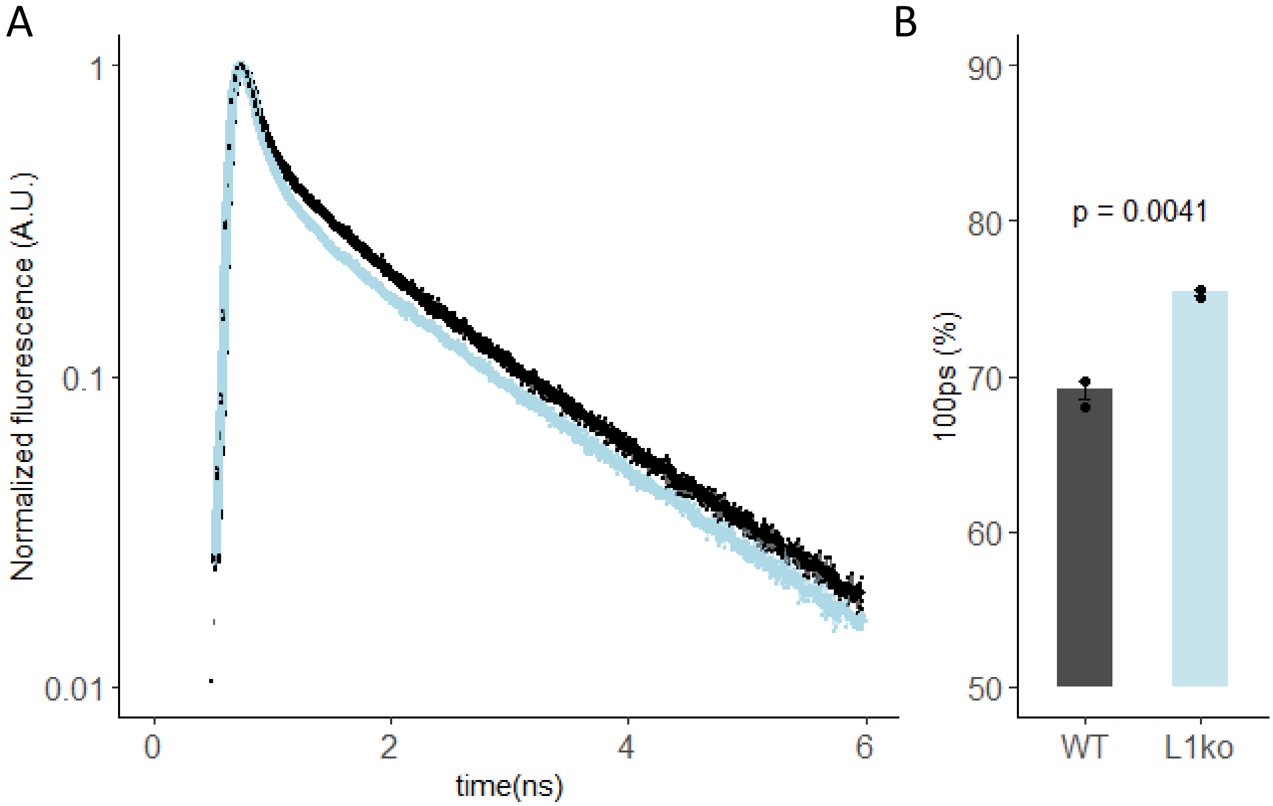


**Figure S6.** Fluorescence lifetime imaging microscopy (FLIM) for L1ko and WT. The FLIM measurement was performed on three individual leaves infiltrated with DCMU from adult plants of WT and L1ko, 116µm*116µm areas were measured. Sample were excited with 633 nm light (40 MHz rep. rate) and fluorescence emission was detected at 710-750 nm. A) Average fluorescence decay traces, with SD, from WT leaves WT (black) and L1ko (light blue). The decay traces were fitted with three exponentials ($\tau_1 = 100ps\ (PSI)$ , $\tau_1 = 900ps$ and $\tau_1 = 2000ps$ (PSII) ). B) Contribution of the fast ($100ps$) decay component in WT in the three measurement. The bar height shows the average contribution for each genotype and the error bar the SE. The 100ps contribution, was used for the calculation of the chlorophyll ratio of PSI/(PSI+PSII) shown in Figure 4 in the main text.

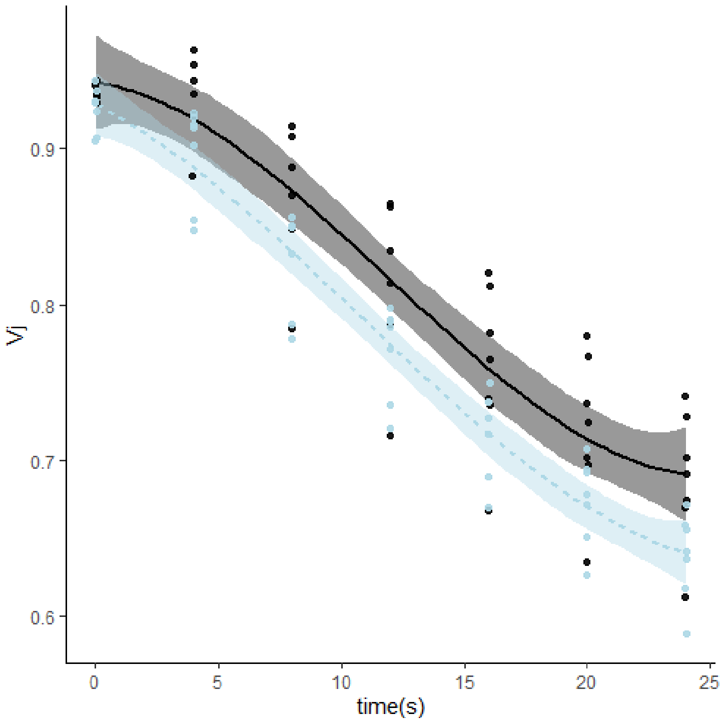


**Figure S7**. Rate of PQ oxidation in dark is similar in L1ko and WT lines. The redox state of the photoactive PQ was inferred from the fluorescence value measured at 3 ms during a 700 ms saturating pulse normalized over the maximal fluorescence ($V_J$), upon sequential incubations of the leaf samples in the dark for increasing time intervals (4-8-12-16-20 and 24 s). The points were fitted with a local regression algorithm based on a second degree function to visually represent the data distribution in function of the time with a continuous error estimate (black continuous line WT, gray line L1ko). The standard error of the interpolation is shown as a gray area around the curves.

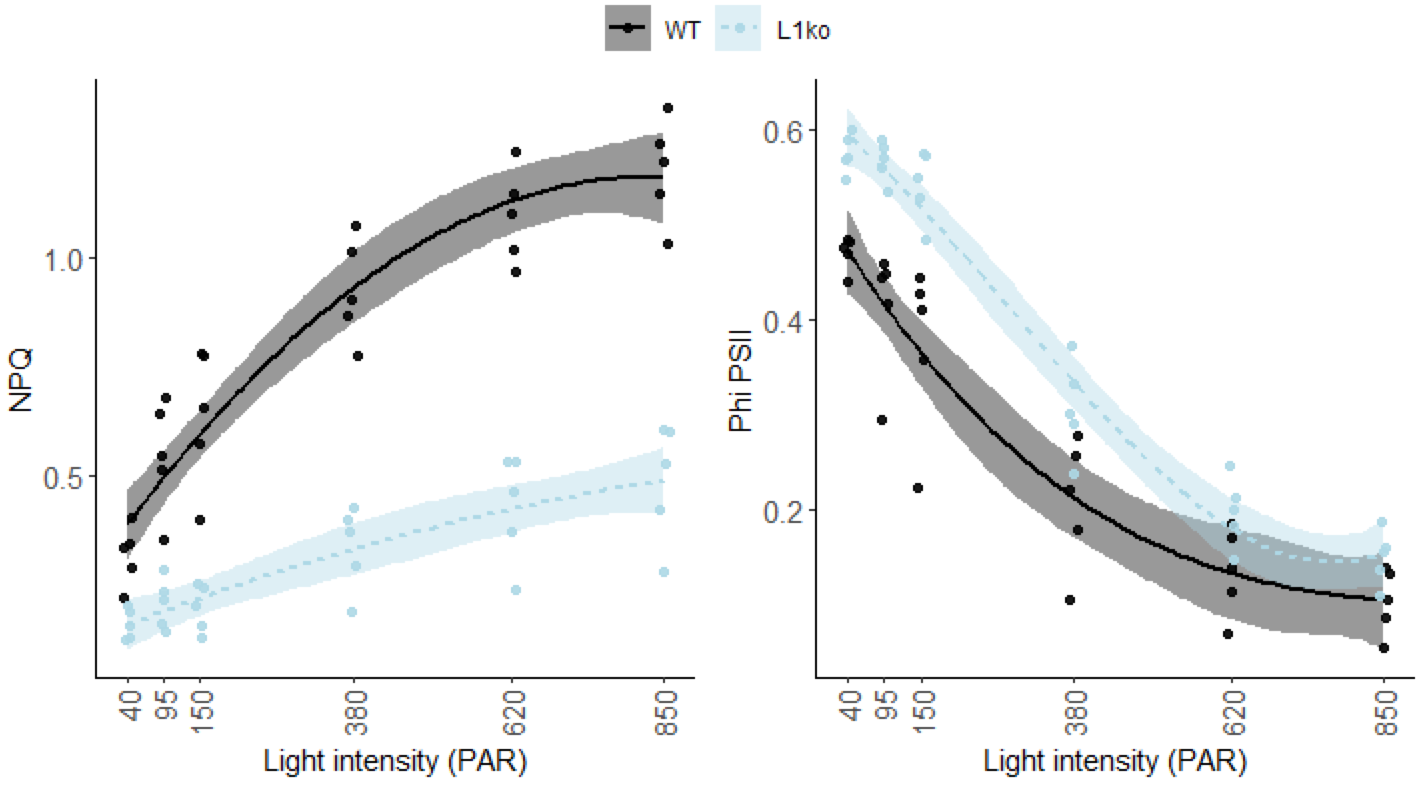


**Figure S8**. Measurement of non photochemical quenching (NPQ) and PSII quantum yield (Phi PSII) with a portable detector. The light curve in figure 4 was repeated with a handheld device (Multispeq v2) on the same 5 different plants. The NPQ and Phi PSII measured for each WT (black dots) and L1ko (Light blue dots) single leaves was plotted in function of the light intensity.
The measured points were fitted with a second degree polynomial function for NPQ, and a third degree polynomial for ϕPSII to visually represent the data distribution and the standard error for WT (Black line and gray area) and L1ko (Blue dotted line and light blue area). Post-hoc analysis of the models for NPQ and ϕPSII show that there is a significant difference between L1ko and WT ($p<0.001$).

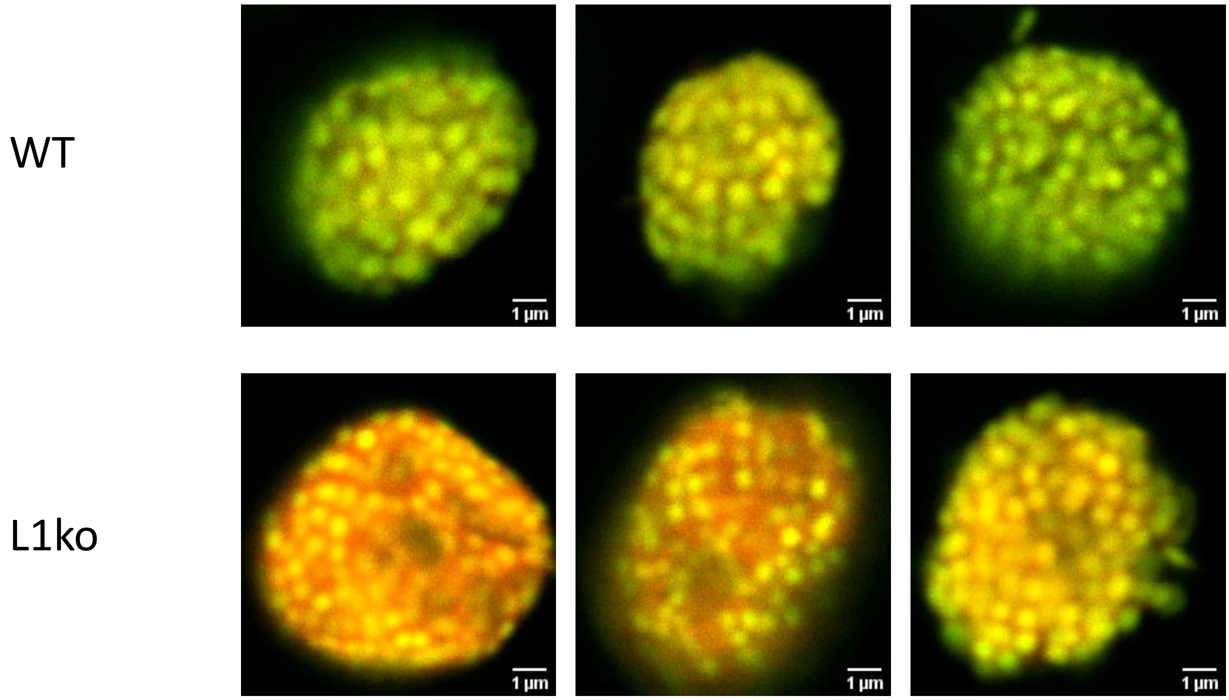


**Figure S9**. Images of the isolated chloroplast from WT and L1ko plants, acquired by confocal microscopy with an excitation at 633nm. The signal at 650-680nm is showed in green (PSII-maximum) while the signal at 710-750nm in red (PSI-maximum). Red channel intensity has been increased 10x in comparison to the green channel to compensate for decreased sensitivity of the detector and decreased fluorescence emission in the far-red region. Bar scale 1µm.

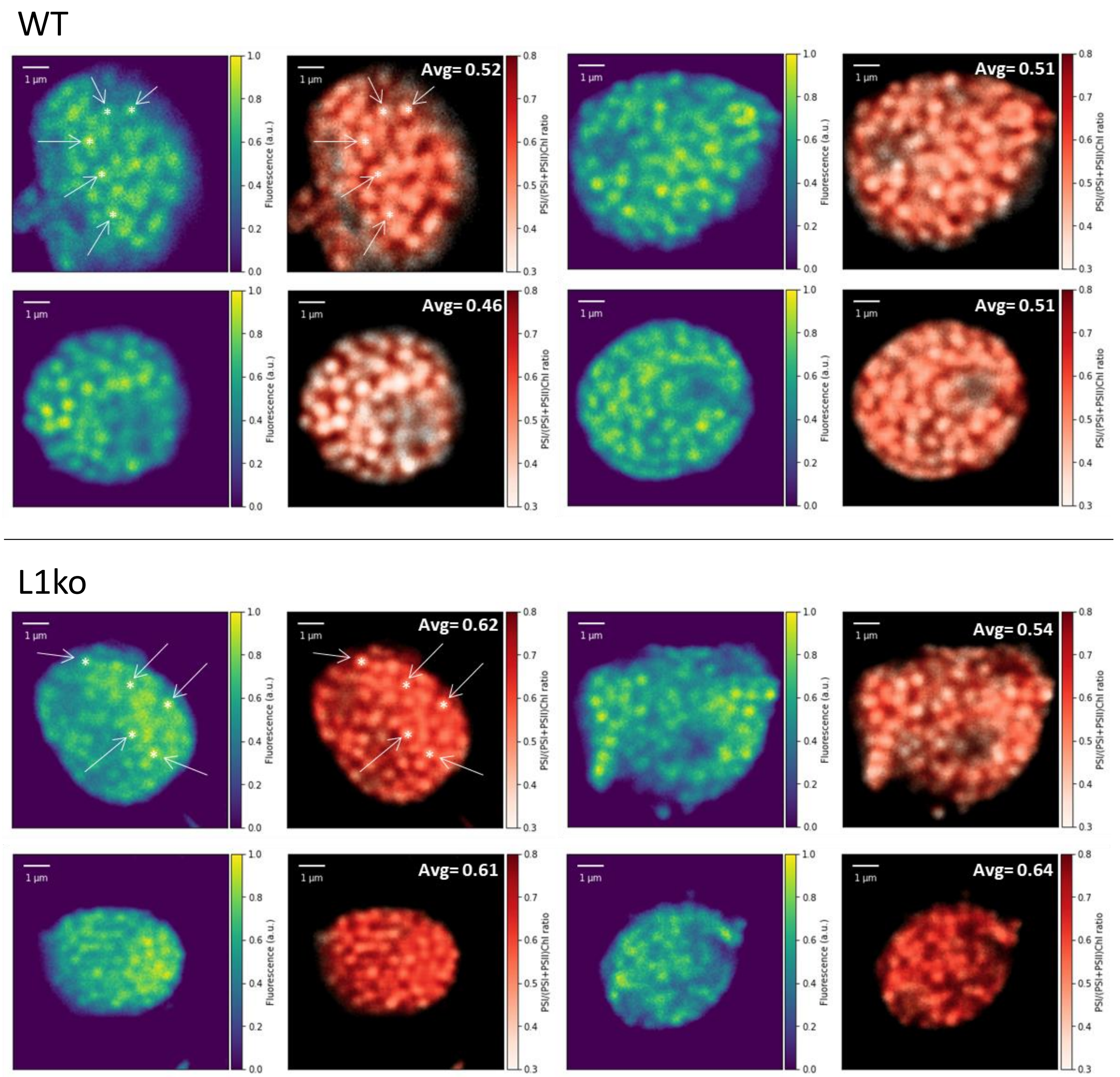


**Figure S10.** Fluorescence intensity and the PSI/(PSI+PSII) Chlorophyll ratio (calculated based on the FLIM measurements according to Wientjes et al. 2017) of isolated chloroplasts from WT and *Lhcb1ko*, incubated with DCMU. The images highlight the grana (lower PSI/(PSI+PSII)Chl ratios) and stromal (higher PSI/(PSI+PSII)Chl ratio) regions. As reference some of the grana are highlighted by an arrow and an asterisk in both the fluorescence intensity image and the PSI/(PSI+PSII) Chl ratio image. The overall PSI/(PSI+PSII)Chl ratio is higher in the WT compared to L1ko and the average ratios calculated, reported in the top right corner, correspond to the measurements made on intact leaves presented in Figure 4. Scale bar 1µm.

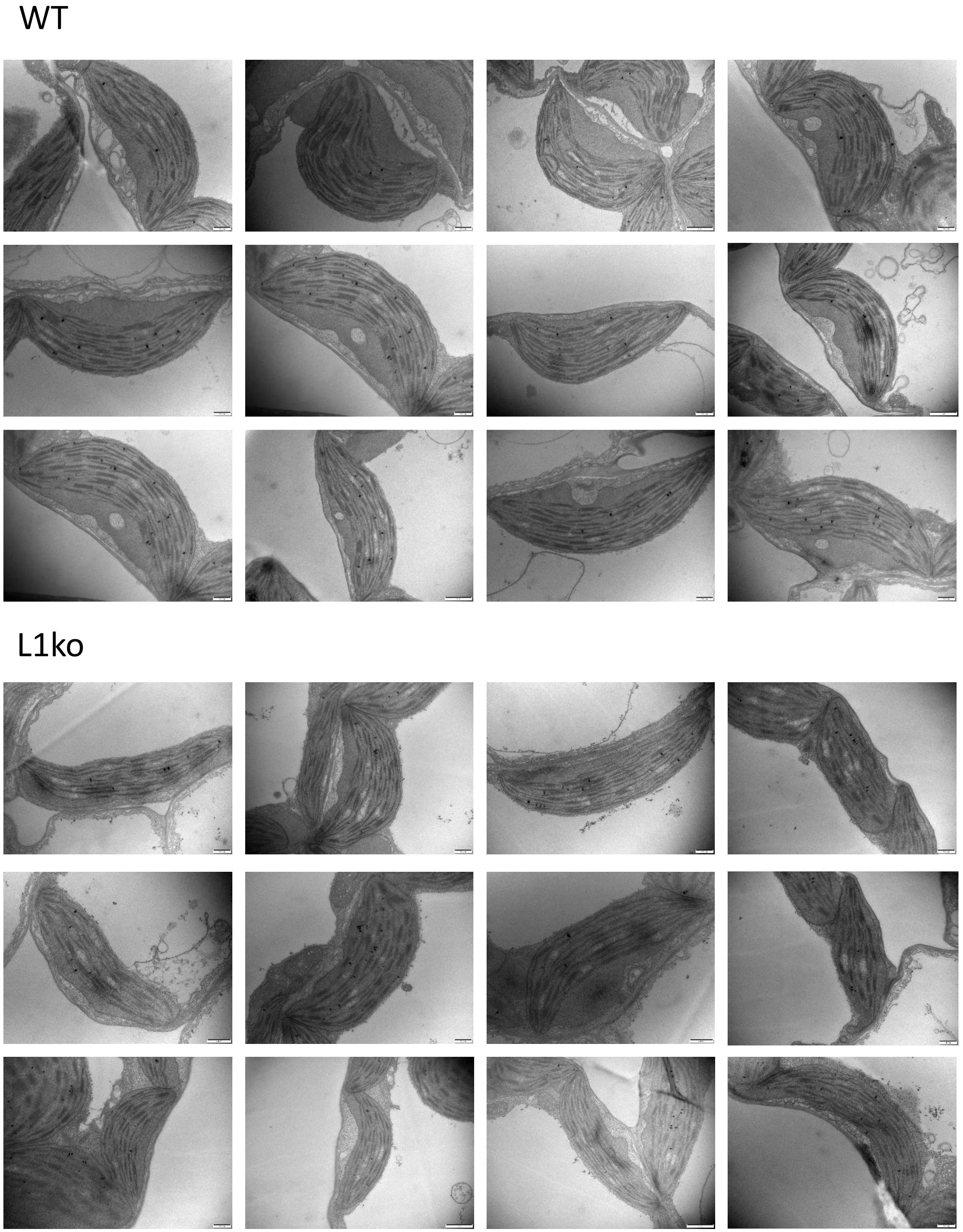


**Supplementary Figure S11.** Collection of TEM images of chloroplasts observed in leaf sections of 15-days old WT and L1ko seedlings. The full set of collected images in the original resolution is available in the Zenodo repository (https://zenodo.org/record/5729177#.YaDZILrjKUk).

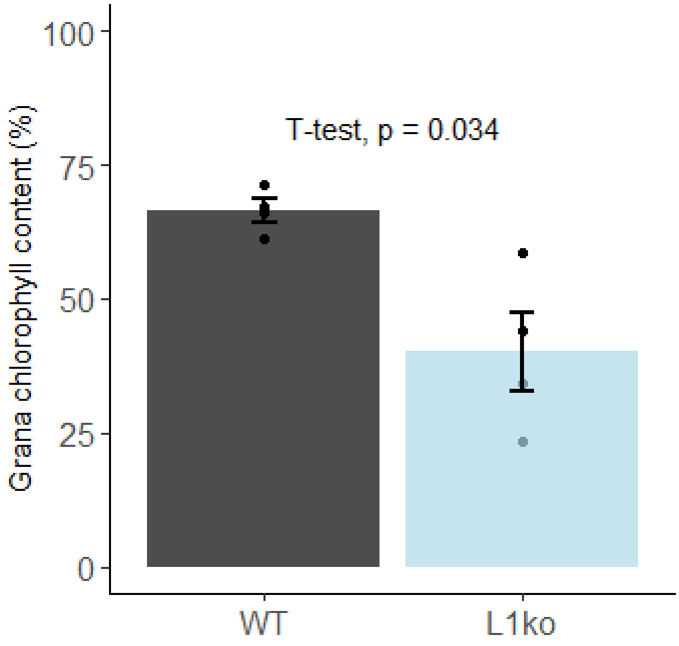


**Figure S12.** The chlorophyll repartition between grana and stroma lameallae is altered in L1ko line. Fraction of the chlorophyll contained in the grana stacks upon fractionation with 1% digitonin, the bar represent the average value for WT (black) and L1ko (Light blue), the error bars represent the standard error, the biological replicates (n=4) are plotted as individual points. The difference was tested by a t-test and the resulting p-value is displayed.

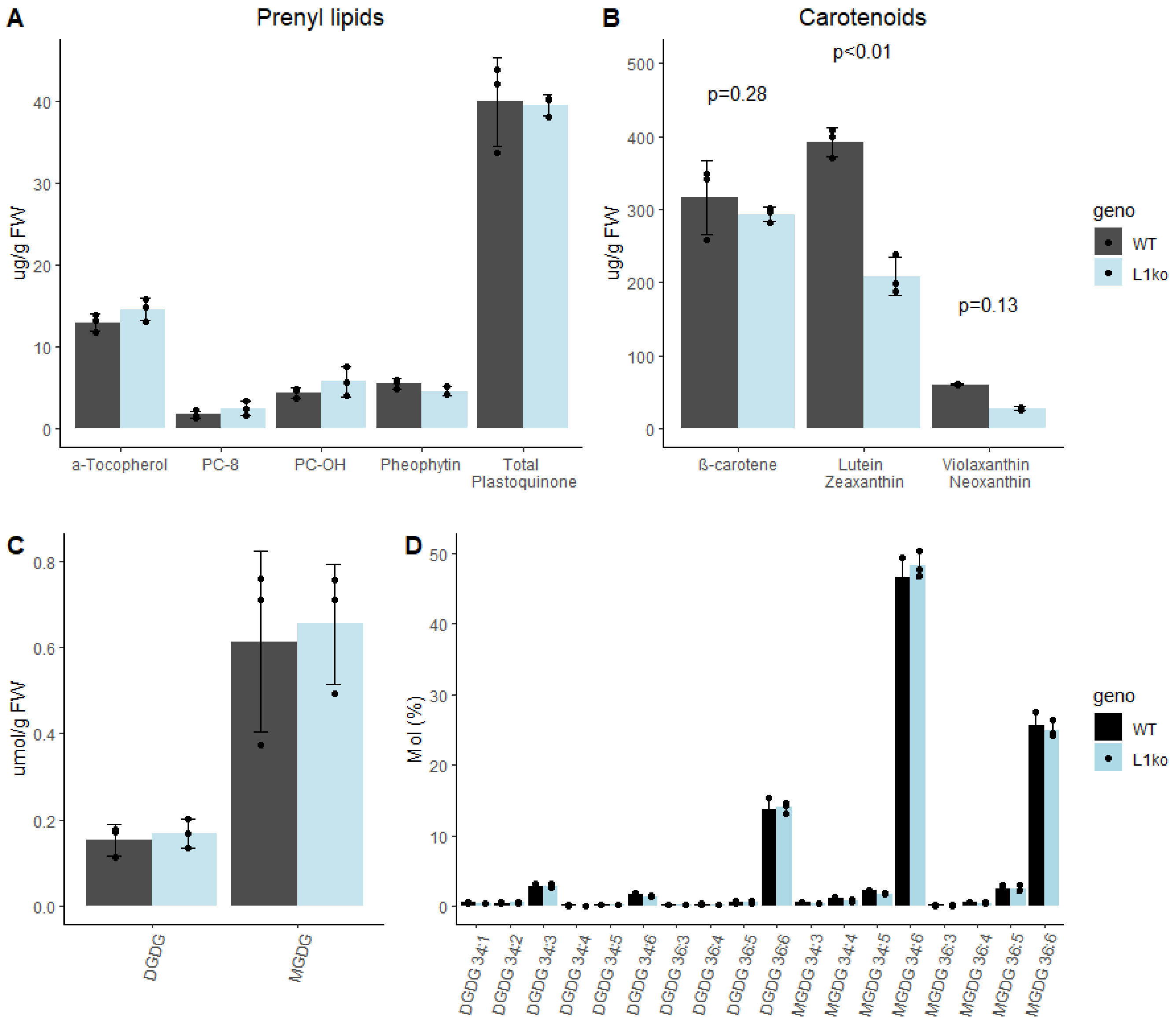


**Figure S13.** Effect of LHCB1 knockout mutation on lipid content. Samples were preparated from WT (Black) and L1ko (Light Blue) adult plants, and the result normalized on the FW used for the extraction. A) Quantification of prenyl-lipids, α -Tocopherol, Plastochromanol (PC-8), Hydroxyl-plastochromanol (PC-OH), Pheophytin and total Plastoquinone. B) Major carotenoids, β-carotene, Lutein and Zeaxanthin, and Violaxanthin with Neoxanthin. C) Galactolipids normalized on fresh weight combined in the classes digalagtosyl-diacyl glycerols (DGDG) and monogalactosyl diacyl glycerols (MGDG). D) detailed profile of the galactolipids divided by the saturation and lenght of the acyl chains. The bars show the average of three biological replicates, the error bars indicate the standard deviation of the mean. The p values in panel B derive from a Student's T-test.
